# Arabidopsis calmodulin-like proteins CML13 and CML14 interact with proteins that have IQ domains

**DOI:** 10.1101/2023.03.09.531943

**Authors:** Howard J. Teresinski, Bryan Hau, Kyle Symonds, Ryan Kilburn, Kim A. Munro, Nathan M. Doner, Robert Mullen, Vivian H. Li, Wayne A. Snedden

## Abstract

In response to Ca^2+^ signals, the evolutionarily-conserved Ca^2+^ sensor calmodulin (CaM) regulates protein targets via direct interaction. Plants possess many CaM-like (CML) proteins, but their binding partners and functions are mostly unknown. Here, using Arabidopsis CML13 as ‘bait’ in a yeast two-hybrid screen, we isolated putative targets from three, unrelated protein families, namely, IQD proteins, calmodulin-binding transcriptional activators (CAMTAs), and myosins, all of which possess tandem isoleucine-glutamine (IQ) structural domains. Using the split-luciferase complementation assay *in planta* and the yeast 2-hybrid system, CML13 and CML14 showed a preference for interaction with tandem over single IQ domains. Relative to CaM, CML13 and CML14 displayed weaker signals when tested with the non-IQ, CaM-binding domain of glutamate decarboxylase or the single IQ domains of CNGC20 (cyclic-nucleotide gated channel-20) or IQM1 (IQ motif protein1). We examined IQD14 as a representative tandem IQ-protein and found that only CaM, CML13, and CML14 interacted with IQD14 among 12 CaM/CMLs tested. CaM, CML13, and CML14 bound *in vitro* to IQD14 in the presence or absence of Ca^2+^. Binding affinities were in the nM range and were higher when two tandem IQ domains from IQD14 were present. Green fluorescent protein-tagged versions of CaM, CML13, and CML14 localized to both the cytosol and nucleus in plant cells but were partially relocalized to the microtubules when co-expressed with IQD14 tagged with mCherry. These and other data are discussed in the context of possible roles for these CMLs in gene regulation via CAMTAs and cytoskeletal activity via myosins and IQD proteins.

## INTRODUCTION

In response to external stimuli and internal cues, plants use a variety of second messengers to process information, with calcium (Ca^2+^) ions being the most prominent and ubiquitous among these. Ca^2+^ signalling is involved in events such as symbioses with rhizobial bacteria and mycorrhizal fungi (Oldroyd & Downie, 2008), response to abiotic (Wilkins et al., 2016) and biotic stress and pathogen attack (Tian et al., 2020; Aldon et al., 2018), as well as during a number of other cellular and developmental events (Dodd et al., 2010; Kudla et al., 2018). Thus, there is an ongoing effort to understand the cellular mechanisms through which Ca^2+^ signals function in plants.

Under resting conditions, cytosolic Ca^2+^ is kept at a low concentration (∼100-200 nM) but is sequestered at high levels (mM) in the apoplast, vacuole and other organelles (Costa et al., 2018; Lee & Seo, 2021). This sequestration facilitates the rapid, passive movement of Ca^2+^ into the cytosol upon stimulus perception, creating localized microdomains and triggering signalling events (McAinsh & Ng, 2013; Mehta & Zhang, 2015; Kudla et al., 2018).

The Ca^2+^ signature hypothesis posits that the complex spatial and temporal patterns of Ca^2+^ influx and efflux, such as waves, spikes and sustained elevation, encode information about the nature of the stimulus to help direct appropriate cellular responses (Tian et al., 2020). The relationship between Ca^2+^ signatures and physiological response is still poorly understood, but recent advances have shown that specific Ca^2+^ signatures are drivers of distinct transcriptional programs, thereby linking Ca^2+^ signals to gene expression (Liu et al., 2015).

Although many of the mechanistic details of the Ca^2+^ signature hypothesis remain unclear, the detection of these signals is tasked to Ca^2+^-binding proteins, termed Ca^2+^ sensors, that reversibly bind Ca^2+^ and facilitate downstream responses via interaction with various targets. In plants, the main Ca^2+^ sensor families include calmodulin (CaM) and CaM-like proteins (CMLs), Ca^2+^-dependent protein kinases (CDPKs), and calcineurin B-like sensors (CBLs) that partner with CBL-interacting protein kinases (CIPKs) (Kudla et al., 2018). As protein kinases, CDPKs and CBL/CIPKs are catalytic responders that detect Ca^2+^ and then post-translationally modify specific substrate targets, whereas CaM and CMLs lack catalytic function and serve as relay regulators through downstream target binding.

CaM is the prototypical Ca^2+^ sensor and is among the most evolutionarily-conserved proteins in eukaryotes (Copley et al., 1999). The Arabidopsis genome encodes seven *CaM* genes representing four isoforms that share 90-100% identity (McCormack & Braam, 2003). Structurally, CaM has been well studied and is an acidic protein of 148 amino acids (17.6 kDa) often described as dumbbell-shaped with N- and C-terminal globular domains joined by a flexible, central helical linker (Ikura & Ames, 2006). The defining structural features of CaM are four Ca^2+^-binding, EF-hand domains, two in each globular region, that reversibly bind Ca^2+^ in a cooperative manner (Kawasaki & Kretsinger, 2017).

When CaM binds Ca^2+^, it undergoes a major conformational change, exposing a large hydrophobic groove that participates in target binding (Zhang et al., 1995). The majority of CaM-target interactions are Ca^2+^ dependent, but there are also targets that bind independently of Ca^2+^, or by a strictly apo-CaM mechanism (Jurado et al., 1999; Houdusse et al., 2006). As a regulatory protein, the physiological roles of CaM are defined by the identity of downstream targets. In plants, the number and diversity of CaM targets is substantial and includes ion channels and pumps, cytoskeletal proteins, metabolic enzymes, transcription factors, and a range of proteins of unknown function (DeFalco et al., 2010). CaM can thus be considered a universal Ca^2+^ sensor in plants and is likely to function in all cells where Ca^2+^ signals occur.

Interestingly, in addition to CaM, plants have evolved a large family of CaM-like proteins, termed CMLs, that are unique to the plant kingdom and some protists (McCormack & Braam, 2003). The seven CaMs and 50 CMLs represented in Arabidopsis are subdivided into nine subgroups, where CMLs range from about 20% to 75% amino acid sequence identity with CaM and, like CaM, possess no functional domains aside from EF-hands. Despite being the largest family of Ca^2+^ sensors in plants, the majority of CMLs are unstudied. However, functions for a few CMLs have been demonstrated (Bender & Snedden, 2013; La Verde et al., 2018), including roles for CML42 in herbivory response (Vadassery et al., 2012) and trichome cell morphology (Dobney et al., 2009), roles for CML37, CML38, and CML39 in biotic and abiotic stress responses (Vanderbeld & Snedden, 2007; Scholz et al., 2014), CML9 in immunity (Leba et al., 2012), and CML24 in autophagy (Tsai et al., 2013). Although most CMLs examined to date show similar biochemical properties to CaM, Arabidopsis CML14 was recently reported to display unusual characteristics (Vallone et al., 2016). CML14, and its close paralog CML13 (95% identical), comprise the sole members of subfamily three and possess ∼50% identity to conserved CaM (McCormack & Braam, 2003). Although the physiological roles of CML13 and CML14 are unknown, CML14 possesses a single functional EF-hand and does not undergo hydrophobic exposure upon Ca^2+^ binding, raising questions about how these properties might affect interaction with targets (Vallone et al., 2016).

A major barrier to understanding the function of plant CMLs has been the lack of identified targets. Evidence suggests that CMLs are unlikely to interact with typical CaM-binding domains (Bender & Snedden, 2013; La Verde et al., 2018) and thus, moving forward, it is important to identify and characterize CML targets in order to delineate the signalling pathways and cellular events that CMLs participate in. Here, we report on the identification of novel putative targets for paralogs CML13 and CML14. These targets include members of the IQ-Domain (IQD), myosin, and CaM-binding transcriptional activator (CAMTA) families. The common structural feature of these protein families is the presence of tandem IQ domains, which are specialized CaM-binding domains defined by the motif IQxxxRGxxxR or the more relaxed (FILV)Qxxx(RK)xxxx(RK) IQ-like motif (Alexander et al., 1988; Bähler & Rhoads, 2002). CAMTAs, myosins, and IQD proteins all possess a minimum of two such IQ domains in close proximity that are arranged in tandem and separated by about 12-20 residues in the primary structure. In plants, IQ domains are also found in cyclic-nucleotide gated channels (CNGCs) and IQ-motif proteins (IQMs), but they occur as single domains, or if present as multiples, are not in a tandem arrangement. IQ domains often bind CaM independently of Ca^2+^ and have mainly been studied in myosins where CaM or CaM-related proteins serve as myosin light chains (MLCs) (Heissler et al. 2014). Among the IQ-targets we identified, we explored in detail the *in planta* and *in vitro* properties of CML13, CML14, and CaM interaction with the IQ domains of a representative IQD family member, IQD14, in order to better understand what roles these CMLs might play in target regulation. Collectively, our data suggest that CML13 and CML14 interact with proteins possessing IQ domains and that they may be unique among CMLs in this respect. We propose a model whereby CML13 and CML14 may complement or substitute for CaM in the regulation of various targets with IQ domains.

## RESULTS

### CML13 & CML14 are highly expressed CML family members

As gene and protein expression databases have expanded considerably since early reports on global *CML* expression (McCormack et al., 2005), we sought to identify Arabidopsis CMLs with high relative expression levels for further analyses. Publicly available proteomic and transcriptomic databases indicate wide diversity in the expression patterns among Arabidopsis *CMLs*. While many exhibit a low basal transcript level, several isoforms, such as *CML6* expression in mature pollen, are specific to an individual developmental stage. Interrogation of proteomic databases drew our attention to CML13 and CML14 as these paralogs were present at high levels and expressed broadly across tissues and developmental stages relative to most CMLs (Supporting Information: Table S1) (Winter et al., 2007; Wang et al., 2015). Developmental transcriptomic data, mined from the BioAnalytical Resource (BAR) at the University of Toronto (Winter et al., 2007) for Arabidopsis *CML13* and *CML14* are presented in Supplemental Information, Figure S1 and Table S2.

To complement and expand upon the BAR transcriptomic data, we sought to analyze the spatiotemporal gene-promoter activity patterns of *CML13* and *CML14*, using β-glucuronidase-(GUS) based *CML* promoter:reporter system in transgenic plants as a qualitative proxy for spatial and developmental patterns of gene expression. Our data suggest that *CML13* and *CML14* have largely overlapping expression patterns (Figure 1), consistent with public transcriptome and proteome database information (Supplemental Figure S1, Supplemental Tables S1, S2). For example, promoters for both isoforms were active in aerial and root tissue, including vascular tissue, in trichomes and siliques, the root elongation zone of seedlings, and around root branch points. In general, the gene expression patterns suggested by GUS reporter analysis indicate that *CML13* and *CML14* are expressed throughout most tissues with the strongest promoter activity in roots, floral organs, and at the base of siliques. The similar expression patterns and high primary sequence conservation suggest that these paralogs likely overlap in function across tissues and developmental stages.

**Figure 1.**
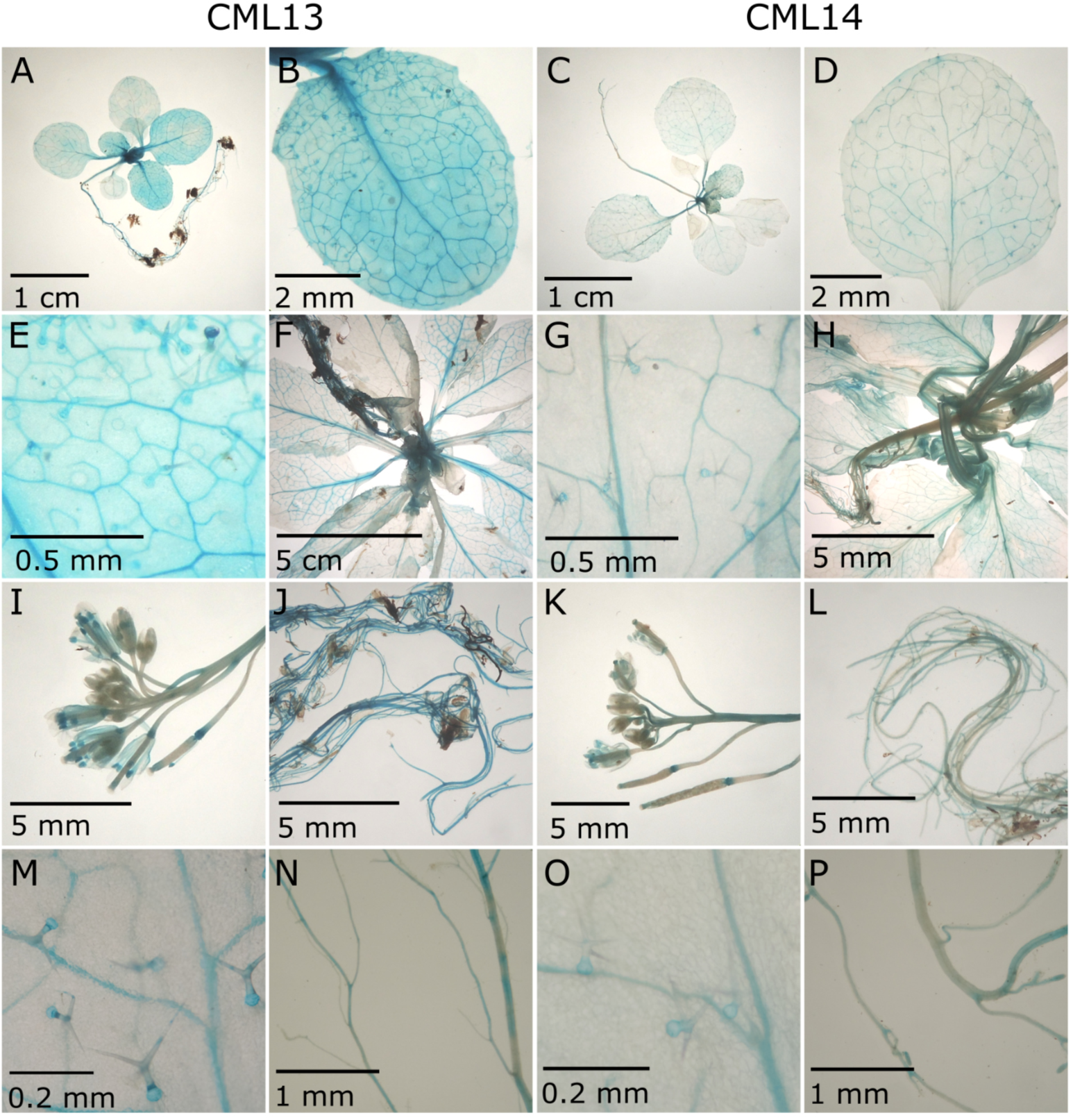
*CML13* and *CML14* promoter activity patterns are broad and similar. Arabidopsis *CML13*pro-GUS and *CML14*pro-GUS transgenic lines were assessed for promoter:reporter activity across various tissues and stages of development. Representative images are shown for (**A, C**) 2-week old seedlings, (**B, D**) 2-week old leaves, (**E, G**) trichomes of 2-week old leaves, (**F, H**) 5-week old rosettes, (**I, K**) 5-week old flowers, (**J, L**) 5-week old roots, (**M**, **O**) closeup of trichomes, (**N**, **P**) closeup of root branches. Arabidopsis were grown on 0.5x MS media (A-H) or on soil (J-L) to the respective growth stage, then fixed and stained as described in Experimental Procedures.

### CML13 and CML14 do not undergo major conformational changes in response to Ca^2+^ binding compared to CaM

CML13 and CML14 possess 95% amino acid sequence identity, differing by only seven amino acids (Figure 2A) making them, aside from CaMs, the two closest relatives in the CML family. Alphafold models (Jumper et al., 2021) predict the tertiary structures of CML13 and CML14 to adopt a partially “closed” conformation, with features reminiscent of both the apo- and Ca^2+^-bound forms of CaM (Figure 2B), consistent with properties of CML14 previously reported (Vallone et al., 2016).

**Figure 2.**
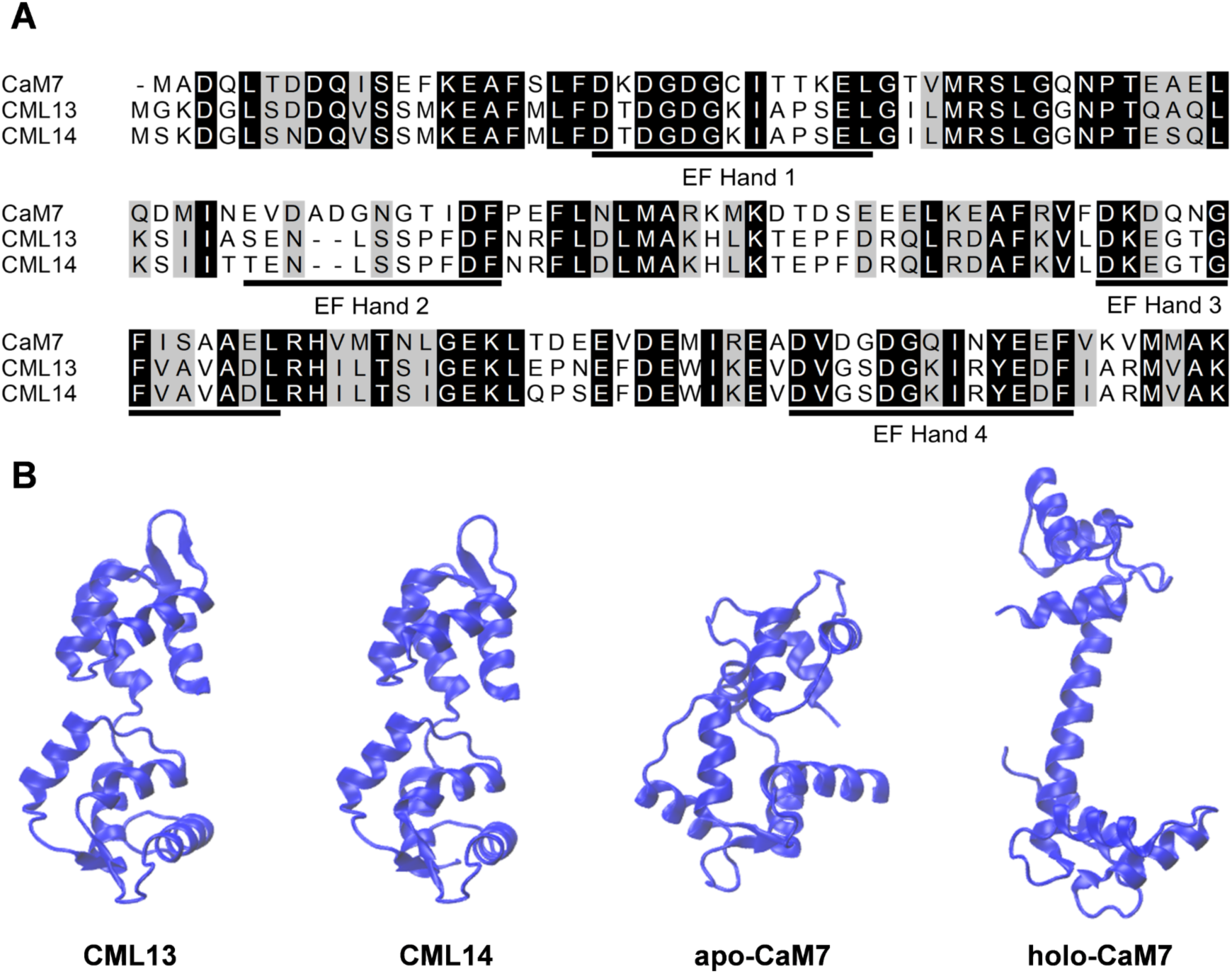
Sequence alignment of Arabidopsis CML13, CML14 and CaM. (**A**) Alignments were performed using ClustalΩ (Sievers and Higgens, 2014). An 85% threshold was used for shading amino acid residues, black if identical, and grey if they possess similar biochemical properties. Predicted and degenerate EF-hands are underlined. CaM7 is a highly conserved CaM isoform presented for comparison. Image generated using Bioedit V7.2.5. (Hall, 1999). (**B**) Predicted ribbon models of CML13, CML14, and CaM in the apo- and Ca^2+^ bound form were generated using the Alphafold protein structure database (Jumper et al., 2021).

Traditional purification strategies for recombinant CaM and CMLs take advantage of their Ca^2+^-induced hydrophobic exposure and the use of hydrophobic-interaction chromatography (Bender et al., 2014). Our early attempts at recombinant expression and purification of CML13 and CML14 revealed that they do not exhibit hydrophobic exposure upon Ca^2+^ binding. These biochemical properties contrast with most other CMLs examined to date (Dobney et al., 2009; Bender et al., 2013; Bender et al., 2014; Ogunrinde et al., 2017) but are consistent with a report on CML14 by Vallone et al., (2016).

We utilized far-UV (180-260 nm) circular dichroism (CD) spectrometry to analyse CML13 and CML14 secondary structures in the presence or absence of Ca^2+^. As expected, CaM exhibited a gain in alpha-helical content in the presence of Ca^2+^, whereas CML13 and CML14 exhibited little to no change (Figure 3A). This extends the findings of Vallone et al., (2016) to include CML13 and indicates that any conformational changes that occur upon Ca^2+^ binding to CML13 or CML14 are not due to major secondary structural rearrangements. In addition, near-UV CD (250-320 nm) was used to detect changes in tertiary structure of CaM, CML13, and CML14 in the presence or absence of Ca^2+^ (Figure 3B). The aromatic residues, Trp, Tyr, Phe, each have distinct spectral profiles in the near-UV range with bands at 290-305 nm, 275-282 nm, and 255-270 nm, respectively (Kelly et al., 2005). CaM, CML13, and CML14 all possess a single, conserved Tyr in the C-term lobe at a position within the fourth EF-hand of CaM (Figure 1). No Trp residues are present in CaM, whereas a single Trp is found in the C-lobe of CML13 and CML14 at position 124 (Figure 2). In contrast, Phe residues, which tend to produce weaker CDS bands, are spread throughout both N-and C-lobes for CaM, CML13, and CML14 (Figure 2). For the near-UV CD spectra, we observed spectral features for each of these aromatics, with the exception of an absent 290-305 nm signal in the Trp region for CaM, as expected. For CML13 and CML14, we did not observe any Ca^2+^-dependent change in tertiary structure fingerprint in any of the three spectral regions (Figure 3B). In sharp contrast, CaM displayed a clear change in the spectral regions associated with both Tyr and Phe. Given that the single Tyr residue in CaM is located in the C-lobe, this indicates that the C-lobe is changing shape in the presence of Ca^2+^. As Phe residues are spread through both C- and N-lobes of CaM (Figure 2), we cannot attribute Ca^2+^-induced changes in the Phe spectral region to either specific lobe. These data thus highlight significant structural similarities between CML13 and CML14 and their differences with CaM with respect to the effects of Ca^2+^ binding.

**Figure 3.**
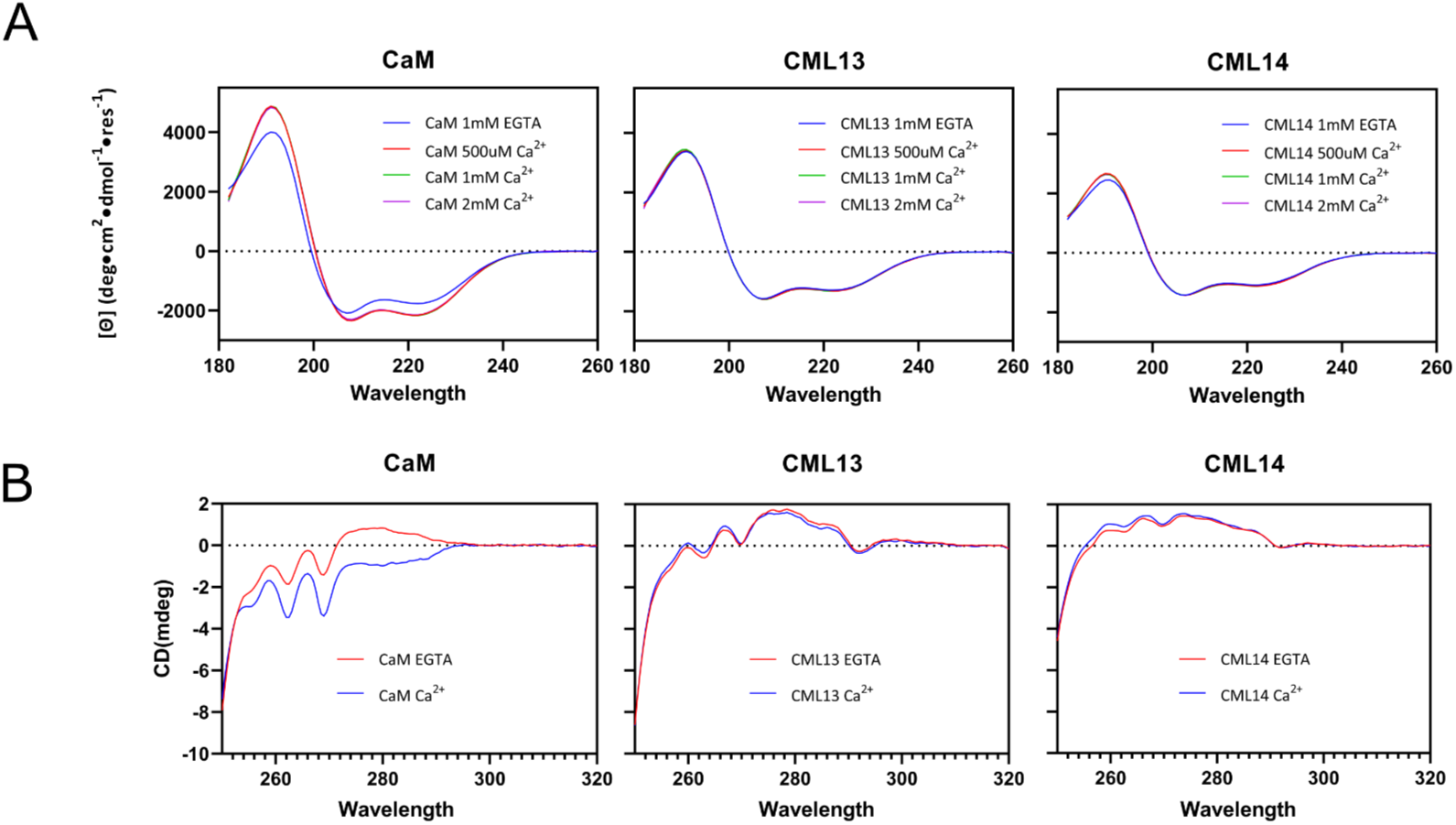
CML13 and CML14 do not display structural changes of typical Ca^2+^ sensors. (**A**) Far-UV CD spectra of pure, recombinant CML13, CML14 and CaM in the presence of 500 μM CaCl2, 1 mM CaCl2, 2 mM CaCl2, or 1 mM EGTA, with corresponding secondary structural composition determined through deconvolution using CONTINLL185. (**B**) Near-UV CD spectra was used to observe tertiary structure changes in CaM, CML13, and CML14 in the presence of 2 mM CaCl2 or 2 mM EGTA. Each plot is a mean of 10 scans. Protein concentrations were 45 μM for CaM and 37 μM for CML13 and CML14.

### Identification of novel binding partners for CML13, CML14, and CaM

Given the unique biochemical properties of CML13 and CML14, we sought to investigate their *in vivo* function by identifying downstream protein targets. To this end, we used CML13 as ‘bait’ to screen an Arabidopsis normalized-cDNA yeast 2-hybrid (Y2H) library.

The Y2H screen identified putative interacting partners for CML13 from three distinct families: IQ67-domain proteins (IQDs), CAMTAs, and myosins. After eliminating false positives that self-activated with an empty bait vector, IQDs, CAMTAs, and a myosin were the only putative interactors isolated in our Y2H screen. In Arabidopsis, these families are represented by 33, 6, and 17 members, respectively. Three IQD family members (IQD13, 14, 26), three CAMTA family members (CAMTA2, 4, 6), and one myosin family member (myosin VIII-B) were isolated in our screen as putative CML13 interactors (Table 1). These interacting proteins were confirmed by independent pairwise re-transformations in the Y2H system and showed no autoactivation of the reporter genes in yeast but strong activation of His and Ade biosynthetic reporters when paired with CML13, indicating a positive interaction (Figure 4A). No other putative targets were isolated in our screen. It is noteworthy that IQDs, CAMTAs, and myosins are functionally and structurally quite distinct with one notable exception; most of the family members of these proteins are predicted to contain multiple, tandem IQ domains (see general domain architecture in Figure 4B). Previous work has established that at least some members from each of these families are CaM targets in plants, but evidence of their interaction with CMLs is very limited (Bouché et al., 2002; Bürstenbinder et al., 2013; Haraguchi et al., 2018). IQ domains generally function as CaM-binding domains, and often bind CaM independently of Ca^2+^ (Andrews et al., 2020). It is also noteworthy that our screen did not identify any Arabidopsis proteins that contain single IQ domains or multiple IQ domains that are not arranged in tandem, such as members of the cyclic-nucleotide gated channel (CNGC) or the IQ-motif (IQM) families, raising the question of whether CML13 and CML14 show a binding preference for tandem rather than isolated IQ domains.

**Table 1.** Putative interacting proteins identified through the CML13 yeast 2-hybrid screen.

| AGI #<sup>1</sup> | TAIR Annotation<sup>2</sup> | Frequency<sup>3</sup> | Predicted # of IQs<sup>4</sup> |
| --- | --- | --- | --- |
| AT3G59690 | IQD 13 | 3 | 2-3 |
| AT2G43680 | IQD 14 | 4 | 2-3 |
| AT3G16490 | IQD 26 | 4 | 2-3 |
| AT5G64220 | Calmodulin-binding transcription activator 2 | 1 | 3 |
| AT1G67310 | Calmodulin-binding transcription activator 4 | 1 | 3 |
| AT3G16940 | Calmodulin-binding transcription activator 6 | 1 | 3 |
| AT4G27370 | Myosin VIII-B | 1 | 3-4 |

<sup>1</sup>Arabidopsis genome initiative annotation of genes.
<sup>2</sup>Common name attributed to the respective genes in TAIR.
<sup>3</sup>Frequency refers to the number of independent colonies in which the prey protein was found in the Y2H screen
<sup>4</sup>Number of sequence specific “IQ” motifs found in each gene

**Figure 4.**
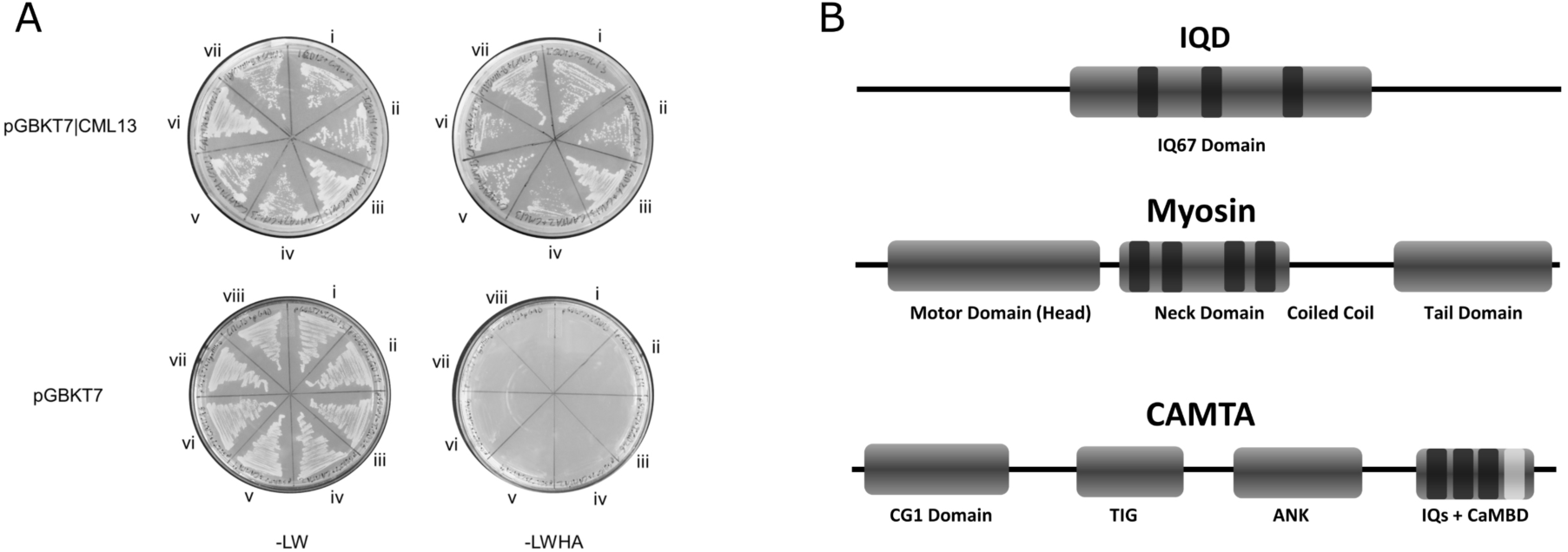
Putative CML13 binding partners tested through pairwise yeast 2-hybrid interaction assays. (**A**) Putative interactors of CML13 that were isolated from a Y2H screen were tested through pairwise retransformation with CML13 bait (upper panels) or empty vector control (lower panels) and the corresponding IQ-domain regions of the respective prey proteins; i, IQD13 (amino acid (AA)# 158-517); ii, IQD14 (AA# 307-668); iii, IQD26 (AA# 98-389); iv, CAMTA2 (AA# 358-1100); v, CAMTA4 (AA# 755-1016); vi, CAMTA6 (AA# 660-845); vii, and myosin VIII-B (AA# 763-1135). Plate section viii denotes yeast-transformed with CML13 bait vector and empty pGADT7 vector as the autoactivation control. Following successful transformation, colonies were restreaked onto -LW (left) or - LWHA (right) media and grown at 30°C for 4 days to verify co-transformation and test for interaction, respectively. Images are representative of at least three independent, pairwise transformations. (**B**) the relative structural architecture of a typical IQD, myosin, and CAMTA protein (TIG, transcription-associated immunoglobulin domain; ANK, ankyrin-repeat domain; CaMBD, CaM-binding domain). Black bands represents predicted IQ domains, and the light-grey band in CAMTA represents a CaM-binding domain (CaMBD).

We analysed the interaction of all putative CML13-binding partners identified in our Y2H screen against CaM and CML14 as baits. Independent, Y2H pairwise-transformations confirmed that CML14 and CaM are also able to interact with the isoforms of myosin, IQDs, and CAMTAs tested in the Y2H system, whereas empty-vector negative controls were not (Figure 4A, Supporting Information: Figure S4).

### Tandem IQ-domain proteins interact with CML13, CML14, and CaM *in planta*

From among the putative targets isolated in our Y2H screen, we selected IQD14, CAMTA4, and myosin VII-B and assessed their *in planta* interaction with CaM, CML13, and CML14 using the split-luciferase complementation system in *Nicotiana benthamiana* (Chen et al., 2008). CML42 was used as a negative control given its low sequence identity to CML13 and CML14 (24%). We used leaf-disc assays and a dedicated luminometer for sensitivity, but representative whole-leaf images are shown in Supplemental Figure S2. These images also highlight the inherent variability in the split-luciferase system, due to uncontrollable differences in bacterial infiltration, infection efficiency, and recombinant protein expression (Bashandy et al., 2015). As presented in Figure 5A, the IQ-regions of IQD14, CAMTA4, and myosin VIII-B showed signals indicative of interaction with CaM, CML13, and CML14, typically ranging from about 3-to 30-fold relative to the negative control (CML42). Immunoblots confirmed expression of CaM/CML bait proteins (Supplemental Figure S3). The statistical tests compare relative, pairwise signals for each CML or CaM bait vs CML42 negative controls with the various IQ-prey proteins. Among the tandem-IQ proteins tested, mean RLU signals for CaM, CML13, CML14 vs CML42 controls were strongest for IQD14 and weakest for CAMTA4, but all showed a statistical difference. Although common practice, it is worth noting that the empty pCAMBIA1300 split-luciferase prey vector does not make an appropriate negative control as it lacks an ATG-start codon in frame with the N-terminal luciferase domain.

**Figure 5.**
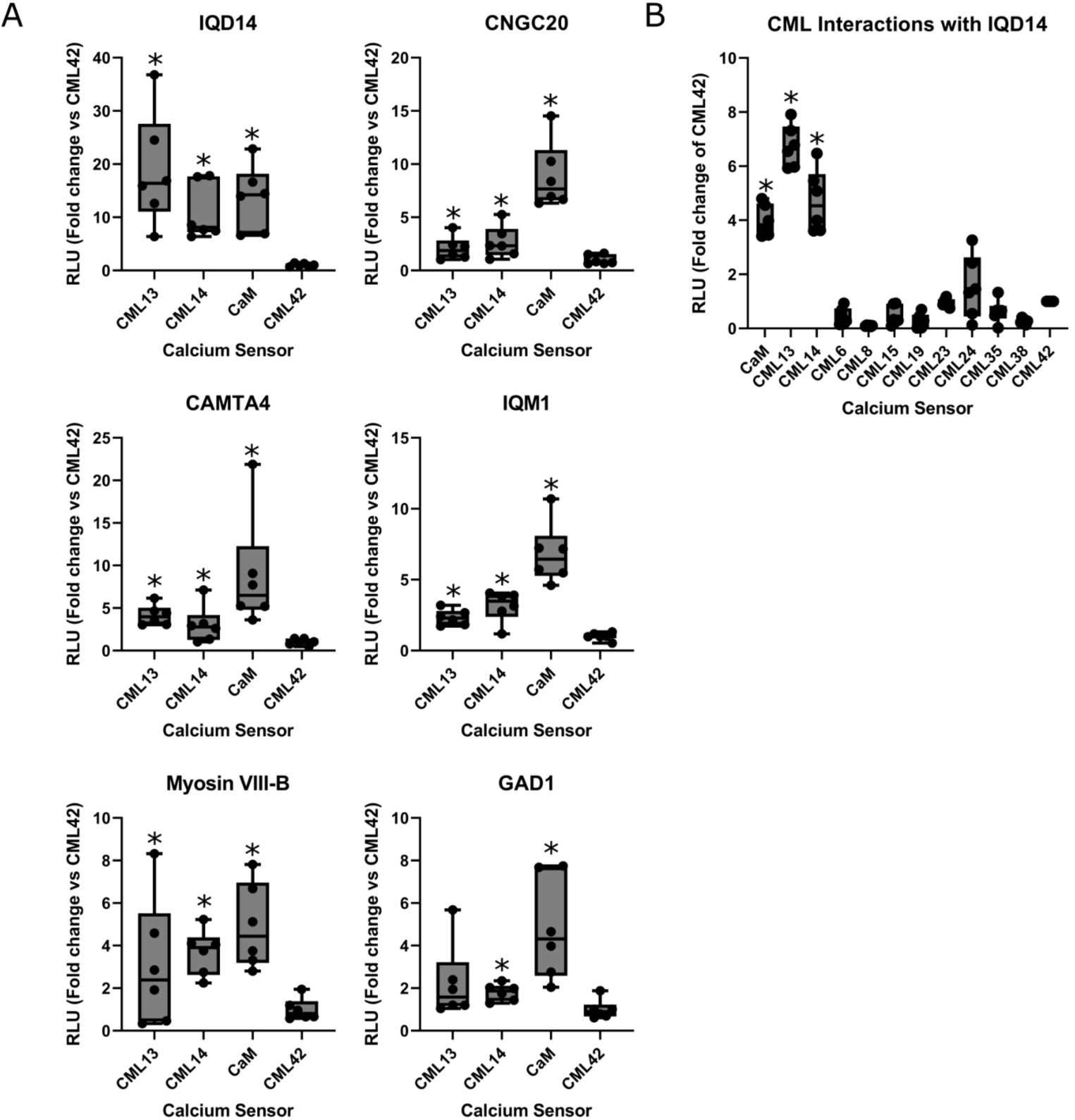
Arabidopsis CaM, CML13, and CML14 interact with representative IQD, CAMTA, myosin family members *in planta*. Box plots of split-luciferase, *in planta* assays to test for (**A**) CML13, CML14, and CaM interaction with representative IQ-domain binding-partner family members or (**B**) the interaction of IQD14 with various CMLs. *N. benthamiana* leaves were infiltrated with *Agrobacterium* harbouring respective NLuc-prey and CLuc-bait (CaM, CMLs) vectors and tested for luciferase activity 3 days later, as described in Experimental Procedures. Data are expressed relative to the RLU signal observed using the negative control bait CML42 which was set to an RLU of 1. Boxes contain each data point for 6 technical replicates, means are shown by a horizontal bar, the grey region is the 95% confidence interval, and whiskers extend to maximum and minimum data points. Asterisks indicate a significantly higher signal vs CLuc-CML42 as a negative control bait (pairwise t-test, p-value < 0.05). Data is representative of at least three independent experiments. RLU; relative light units.

Given that our Y2H screen did not isolate targets with only CaM-binding or single IQ domains, we explored whether CML13 and CML14 could efficiently bind to the non-IQ, CaM-binding domain of Arabidopsis glutamate decarboxylase (AtGAD1), or the single-IQ domain of IQM1 or CNGC20, using CaM as a positive control. As predicted, mean relative light values for CaM were higher, but CML13 and CML14 mean values were typically greater than the negative control, suggesting some level of interaction above background with these non-IQ CaM-binding domains (Figure 5A). To further assess the specificity of CML interaction with IQ domains, we tested nine additional CMLs, drawn from subgroups across the CML family, with the IQ-region of IQD14 as a representative multi-IQ prey in the split-luciferase system. As shown (Figure 5B, Supplemental Figure 2), among the 12 CaM/CMLs tested, only CaM, CML13, and CML14 exhibited a significant signal relative to the CML42 negative control, indicating clear specificity of interaction among CMLs. Collectively, these data suggest that CaM, CML13 and CML14 are the able to bind to IQ-domains whereas most CMLs are not.

### GFP-fusions of CaM, CML13, and CML14 co-localize with IQD14-mCherry at microtubules in plant cells

We next sought to determine the subcellular localization of CML13 and CML14 and further test their interaction with the IQ-region of IQD14 in plant cells. To that end, we generated fusion constructs consisting of two copies of the green fluorescent protein (GFPx2) linked to CML13 or CML14, as well as to CaM. Each fusion protein was then transiently-coexpressed with the nuclear marker protein NLS-RFP consisting of nuclear localization signal linked to three copies of the red fluorescent protein (Dhanoa et al., 2010) in *N. tabacum* Bright Yellow-2 (BY-2) suspension-cultured cells, which are a well-known model system for studying plant protein localization *in vivo* (Brandizzi et al., 2003; Miao & Jiang, 2007). As shown in the epifluorescence micrographs in Figure 6A, and consistent with the majority of other CMLs examined to date (Inzé et al., 2012; Vadassery et al., 2012; Bender et al., 2014), CaM-GFPx2, CML13-GFPx2 and CML14-GFPx2 localized to both the cytoplasm and nucleus (i.e., nucleoplasm) in BY-2 cells. Notably, GFPx2 alone also localized to the cytoplasm and nucleus (Figure 6A).

**Figure 6.**
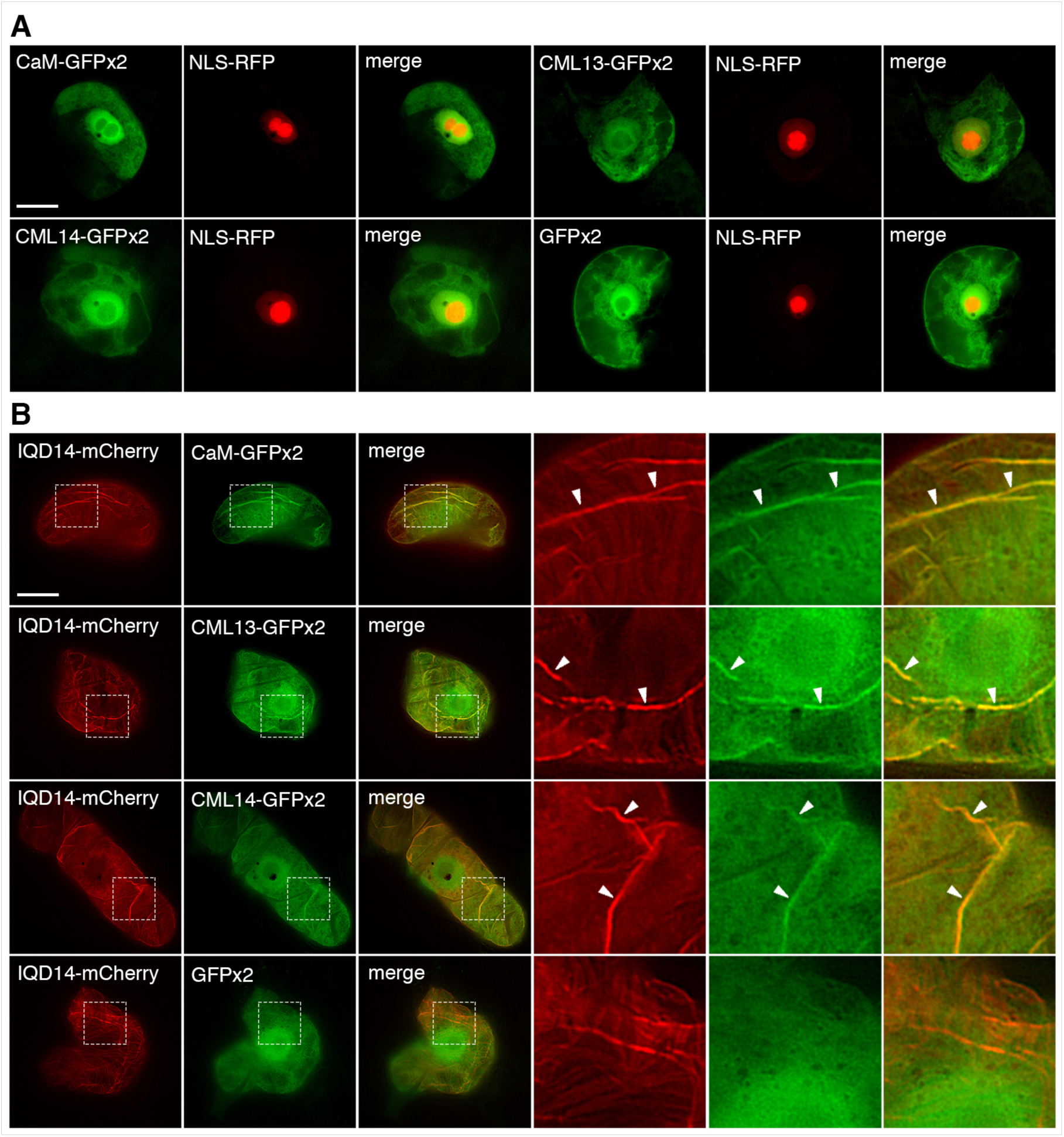
CaM, CML13, and CML14 are partially relocalized to microtubules in BY-2 cells co-expressing IQD14. (**A**) Subcellular localization of GFPx2 fusions of CaM, CML13 and CML14 in BY-2 cells. Cells were transiently-cotransformed (as indicated by labels) with either CaM-GFPx2, CML13-GFPx2 or CML14-GFPx2 (or GFPx2 alone) and NLS-RFP, serving as a nuclear marker protein, and then subsequently incubated for ∼5 hours, formaldehyde-fixed, and imaged using epi-fluorescence microscopy. Shown are representative images, as well as the corresponding merged images for each set of co-transformed cells. Note the localization of each GFPx2 fusion protein throughout the cytosol and nucleus (nucleoplasm) and the accumulation of NLS-RFP primarily in the nucleolus. Bar = 10 μm. (**B**) Subcellular localization of GFPx2 fusions of CaM, CML13 or CML 14 (or GFPx2 alone) in cells co-expressing the IDQ14-mCherry, as indicated by labels. Shown are representative images, along with the corresponding merged images; boxes represent the portion of the cells shown at higher magnification in the panels to the right. Solid arrowheads indicate obvious examples of the CaM/CML13/CML14-GFPx2 fusion protein colocalized with IQD14^I^-mCherry at microtubules. By contrast, note the lack of colocalization of GFPx2 alone with IQD14-mCherry at microtubules. Bar = 10 μm.

Given the results from a previous study showing the subcellular re-localization of CaM when co-expressed with various IQD family proteins (Bürstenbinder et al., 2017), we selected IQD14 for further exploration of *in vivo* interaction with CML13 and CML14, using CaM as a positive control. Consistent with its previously reported localization in plant cells (Bürstenbinder et al., 2017), full-length IQD14 localized to microtubules in BY-2 cells (Figure 6B). Co-expression of CML13-GFPx2, CML14-GFPx2, or CaM-GFPx2 with IQD14-mCherry, but not GFPx2 alone co-expressed with IQD14-mCherry, resulted in their partial re-localization to microtubules (compare Figure 6B with images of CaM/CML13/CML14-GFPx2 in Figure 6A). These data support our Y2H and *in planta* split-luciferase protein-protein interaction analyses and suggest that IQD14 can recruit CML13 and CML14 to the cytoskeleton.

### Delineation of the interaction regions of IQD14 with CaM, CML13, and CML14

We performed a series of truncations with IQD14 as a representative target in the pairwise Y2H transformation system to delineate the interaction region with CML13, CML14, and CaM, while CML42 and empty pGADT7 were used as negative controls. CML13 and CML14 showed identical interaction profiles and were able to bind with any combination of multiple IQ domains but not with all single IQ domain constructs (Figure 7). CaM showed a interaction profile similar to that of the CMLs with the exception that it did not interact with the IQ2+3 combination of IQD14 and interacted with any region containing the first IQ domain. CNGC20, a known CaM-interacting protein with a single IQ domain (Fischer et al., 2013), was used as a positive control for CaM. Consistent with our split-luciferase assay (Figure 5A), CaM interacted well with CNGC20 as expected (Supplemental Figure S4), whereas CML13 and CML14 did not, further supporting the hypothesis that CML13 and CML14 display a preference for specific or multiple IQ domains for efficient binding *in vivo*.

**Figure 7.**
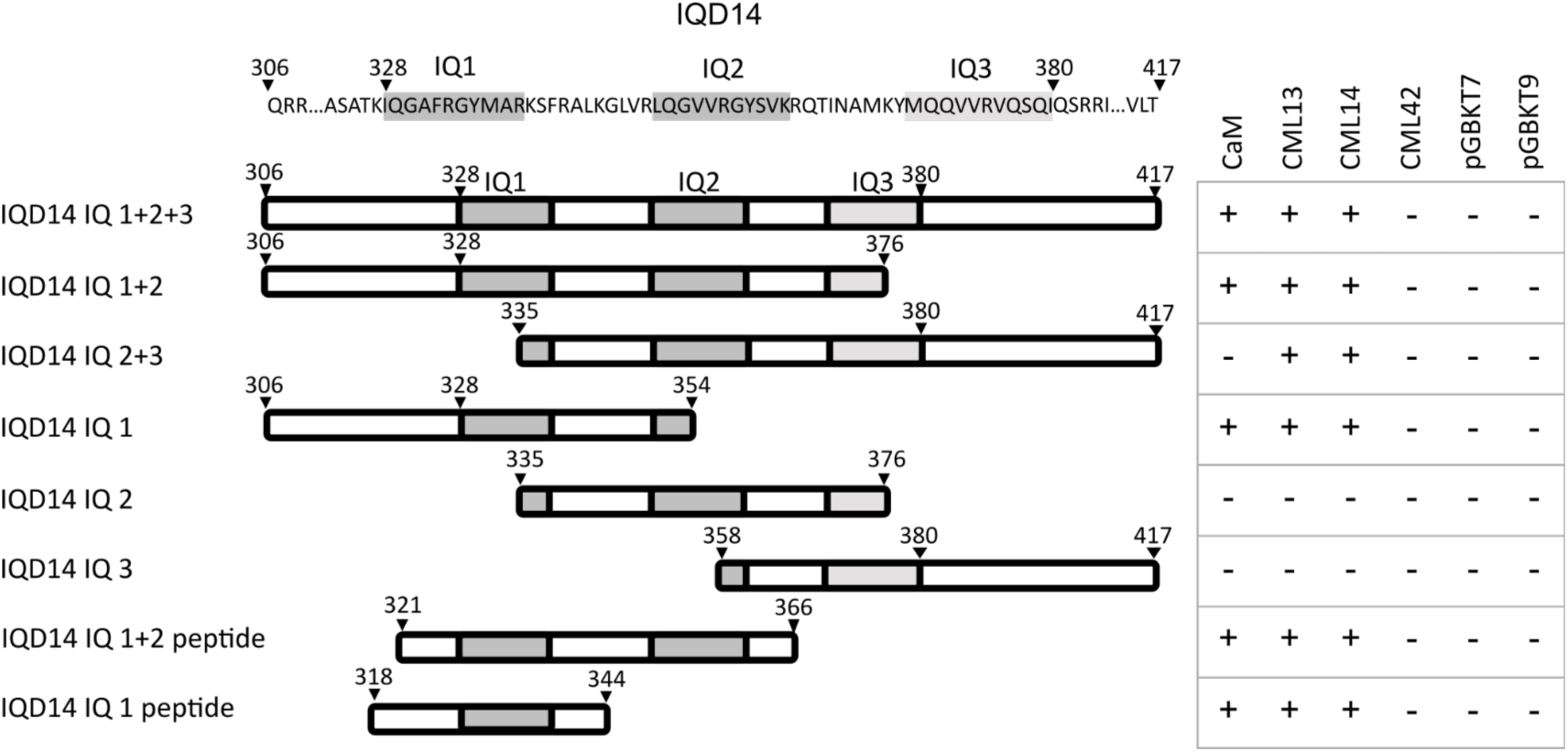
CaM, CML13, and CML14 interact with specific IQ domains in IQD14. Pairwise yeast two-hybrid assays were performed using various delineations of the IQ-domain region of IQD14. A schematic representation of the delineation constructs is presented with the corresponding amino acid sequence and position. IQ motifs are shaded in dark grey and the degenerate IQ motif (IQ3) is shaded in light grey. Bait proteins used in co-transformations are listed in columns in the right panel, with growth on -LWH media being denoted as a “+” and “-” for positive or negative interaction, respectively. For images of pairwise transformation data, see supplemental figure S4.

### Analysis of CML13, CML14, and CaM interaction with the IQ domains of IQD14

In order to further characterize the interactions of IQ domains with CaM and CML13 and CML14 under more controlled conditions, we performed various *in vitro* biophysical analyses using recombinant proteins corresponding to the IQ-domain region of IQD14. As previously documented (Homma et al., 2000; Bürstenbinder et al., 2017), we found that recombinant proteins with multiple IQs were insoluble, regardless of the *E. coli* strain, purification tag, or growth condition used. However, we were able to perform spot-blot overlay assays using purified, recombinant IQD14 containing all three putative IQ domains solubilized in 8 M urea. We covalently conjugated biotin to CaM, CML13 and CML14 and used strepavidin-HRP to detect their interaction with IQD14 or BSA (negative control) spotted onto nitrocellulose. CaM, CML13, and CML14 all bound specifically to IQD14-IQ1+2+3 in the presence or absence of Ca^2+^, but not to BSA negative controls (Figure 8). Additional control overlay assays testing CML42 *in vitro* binding to IQD14 showed very weak interaction in comparison to CaM as a positive control (Supplemental Information: Figure S6).

**Figure 8.**
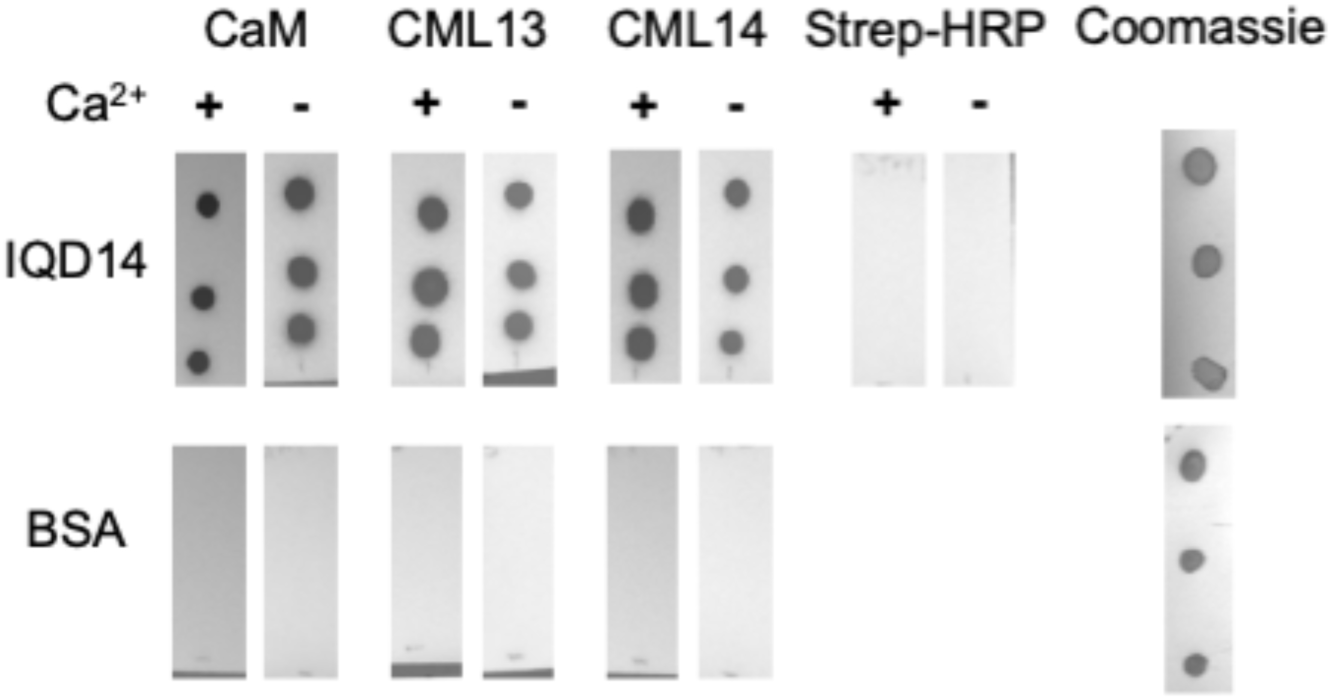
*In vitro* protein-interaction overlay assays show interaction of CaM, CML13, CML14 with IQD14. Triplicate samples of pure, recombinant IQD14 (upper panels, 200 ng, residues 307-418) or BSA (200 ng, lower panels) as a negative control, were spotted onto nitrocellulose, blocked with 5% casein, and incubated with 200 nM biotinylated CaM, CML13, or CML14, as indicated, in the presence of 2 mM CaCl2 (“+”) or 5 mM EGTA (“-”) and interaction was detected using streptavidin-HRP and chemiluminescence. The Strep-HRP (negative control) panels indicate that IQD14 does not interact with streptavidin-HRP alone. Representative Coomassie-stained blots are presented on the far right. Data are representative of a minimum of three independent experiments.

### Biophysical analyses of CaM, CML13, and CML14 interaction with IQ domains of IQD14

We used ITC, in conjunction with synthetic peptides corresponding to the IQ-domain region within IQD14 (see Figure 9A for peptide sequences), to explore the binding of recombinant CaM, CML13, and CML14 bind to the IQ domains and to elucidate the biophysical properties of those binding events. As the third putative IQ domain in IQD14 (IQ3) is very poorly conserved and is not predicted by most motif-searching software, we restricted our analysis to the two conserved IQ domains, IQ1+2. Representative ITC plots for CaM and CML14 are presented in Figure 9 and plots for CML13 are found in the Supplemental Information, Figure S5. Binding affinity data are summarised in Table 2 and thermodynamic values are summarized in Supplementary Information: Tables S4, S5. Interaction of the IQD14-IQ1+2 peptide with CaM, CML13, and CML14 occurred through a series of binding events. In the presence of EGTA, binding of each of the three Ca^2+^ sensors appears to be similar in mechanism, albeit with variable affinities; one high-affinity binding event followed by another weaker binding event. The binding of CML13 and CML14 to IQD14-IQ1+2 in the presence of Ca^2+^ was similar, however, CML13 displayed a more notable decrease in interaction affinities with Ca^2+^ present compared to CML14 (Table 2, Supplemental Information: Figure S5). Binding of CaM to IQD14-IQ1+2 in the presence of Ca^2+^ vs EGTA seems to occur via a different mechanism, although affinities under both conditions were comparable.

**Figure 9.**
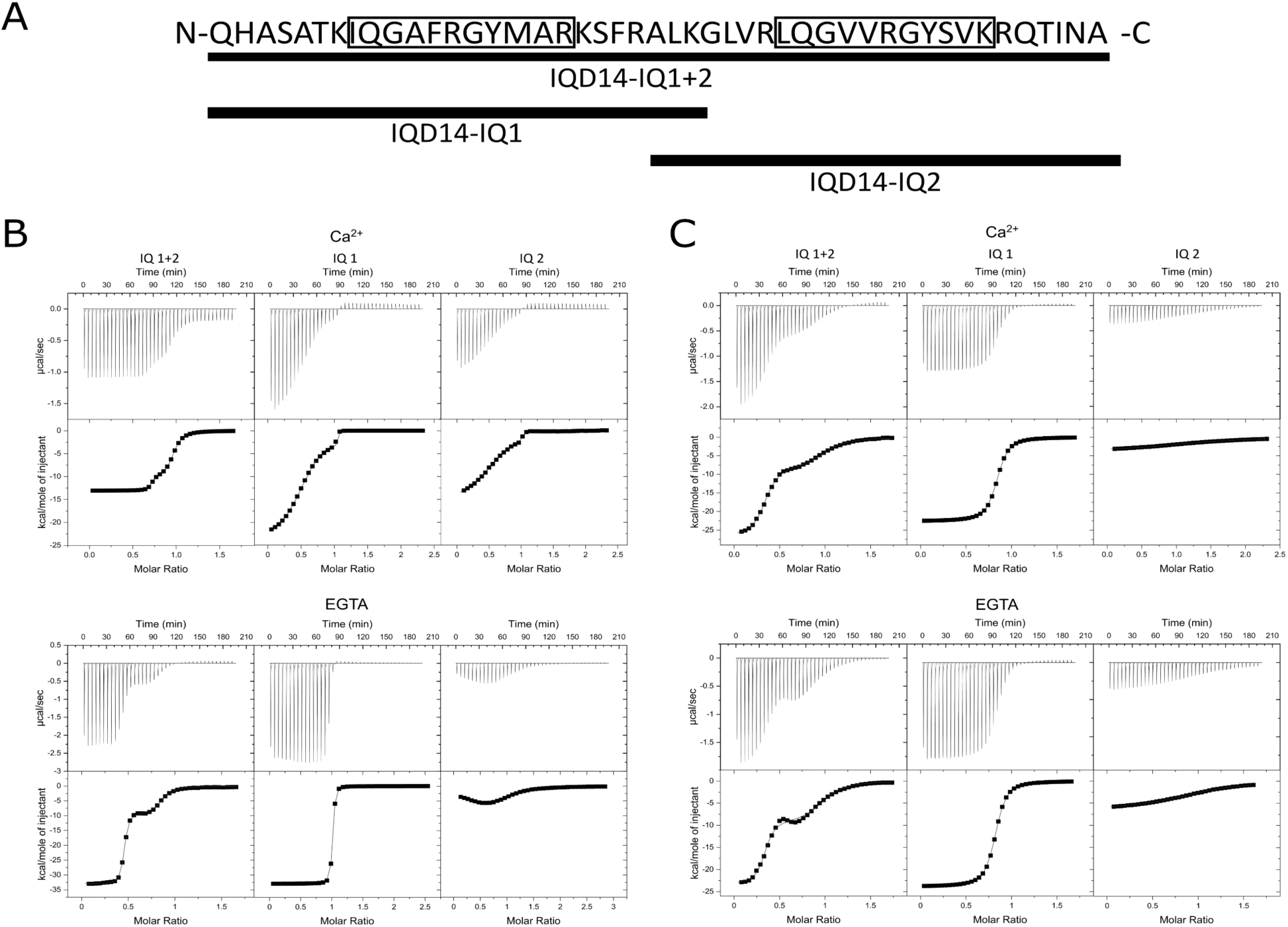
CaM and CML14 display different binding properties to IQ domains of IQD14. (**A**) Region of IQD14 (residues 322-367), with the IQ domains boxed, tested for binding using ITC analysis. The underlined regions correspond to the synthetic peptides used in binding assays. Representative thermogram plots for CaM (**B**) and CML14 (**C**) are shown. Summarized binding data for CML13, CML14, and CaM are presented in Table 2. (**B**) Recombinant CaM (B, left to right; 252 μM , 204 μM, 204 μM) was titrated into IQD14-IQ1+2 (left; 34.3 μM and 39 μM, in EGTA and CaCl2 respectively), IQD14-IQ1(center; 28.8 μM), and IQD14-IQ2 (right; 30 μM) in the presence of 2 mM CaCl2 (top) or 2 mM EGTA (bottom) with 7-μL injections at 300-second intervals at 30°C. (**C**) Recombinant CML14 (252, 225, 430 μM, left to right in CaCl2 and 494, 311 and 311 μM, left to right, in EGTA) was titrated into IQD14-IQ1+2 (39 μM), IQD14-IQ1 (30 μM and, 28.8 μM in Ca^2+^ and EGTA respectively), and IQD14-IQ2 (42 μM and 30 μM in Ca2+ and EGTA respectively) in 7-μL injections at 300-second intervals and 30°C in 10 mM Tris-HCl pH 7.5, 2 mM EGTA. Resulting thermographs were fitted using OriginPro and the appropriate one- or two-site binding model and are representative of at least three independent experiments.

**Table 2.** Summary of interaction affinities obtained by isothermal titration calorimetry (ITC) for CaM, CML13, and CML14 binding to IQD14 peptides.

| Titrant<sup>1</sup> | Peptide<sup>2</sup> | Ca<sup>2+</sup> |  | EGTA |  |
| --- | --- | --- | --- | --- | --- |
|  |  | K<sub>D</sub><sup>3</sup> | n<sup>4</sup> | K<sub>D</sub> | n |
| CaM | IQ1+2 | 1.18 ± 0.17 nM | 0.695 | 1.00 ± 0.16 nM | 0.438 |
|  |  | 364 ± 14.1 nM | 0.260 | 379 ± 57.1 nM | 0.415 |
|  | IQ1 | 0.92 ± 0.63 nM | 0.329 | 7.00 ± 0.70 nM | 0.985 |
|  |  | 8.62 ± 6.47 nM | 0.378 |  |  |
|  | IQ2 | 3.21 ± 1.26 nM | 0.479 | 209 ± 19.4 nM | 0.240 |
|  |  | 21.90 ± 9.91 nM | 0.513 | 1.32 ± 0.03 μM | 0.377 |
| CML13 | IQ1+2 | 217 ± 25.6 nM | 0.348 | 3.48 ± 1.96 nM | 0.316 |
|  |  | 16.20 ± 1.81 μM | 0.725 | 344 ± 200 nM | 0.881 |
|  | IQ1 | 935 ± 16.8 nM | 0.801 | 222 ± 9.19 nM | 0.999 |
|  | IQ2 | 57.50 ± 1.62 μM | 1.16 | 39.7 ± 1.19 μM | 0.981 |
| CML14 | IQ1+2 | 27.5 ± 2.46 nM | 0.328 | 11.50 ± 2.75 nM | 0.323 |
|  |  | 1.11 ± 0.10 μM | 0.668 | 893 ± 175 nM | 0.657 |
|  | IQ1 | 242 ± 3.17 nM | 0.825 | 116 ± 2.77 nM | 0.810 |
|  | IQ2 | 5.24 ± 0.07 μM | 0.882 | 3.16 ± 0.05 μM | 0.996 |

<sup>1</sup>Recombinant proteins were titrated into the ITC cell for each experiment.
<sup>2</sup>Synthetic IQD14 peptides as described in Experimental Procedures.
<sup>3</sup>K<sub>D</sub> was calculated using Originlab2020 ITC fitting software. Values are mean ± SE
<sup>4</sup>n is stoichiometry of binding event. If multiple rows are present, it represents multiple binding events.

In general, the binding curves CaM/CMLs to IQD14 peptides containing a single IQ domain were less complex than those observed using the IQD14-IQ1+2 peptide with two IQ domains. Binding of CaM or CMLs to the IQ1 peptide in EGTA occurred as a single event, with CaM having a higher affinity (*K*D values of 7.00 nM for CaM, compared to 222 nM and 116 nM for CML13 and CML14, respectively). As noted in the analyses with the IQD14-IQ1+2 peptide, CML14 showed a minor decrease in interaction affinity in the presence of Ca^2+^, whereas CML13 again displayed a more substantial decrease. Interestingly, CaM appears to have a different binding mechanism to single IQ domains in the presence of Ca^2+^. Rather than one binding event as seen in the presence of EGTA, we observed two high-affinity binding events (*K*D values of 0.92 nM and 8.62 nM). It is noteworthy that CML13 and CML14 were found to bind IQ1+2 peptide with nM affinity but bound the single-IQ2 peptide weakly, with micromolar affinities that are unlikely to be physiologically relevant, whereas CaM bound IQ2 with properties similar to that observed for IQ1 in Ca^2+^ (ie. two high-affinity binding events). Moreover, in comparison to other treatments, CaM bound the IQ2 peptide in a distinct manner in the presence of Ca^2+^.

Collectively, the ITC data suggest that CaM and CML13,14 interactions occur with physiologically-relevant (nM) affinities when tandem IQ domains are present, these events are mechanistically complex, and there appear to be subtle differences in the way CML13 and CML14 bind to IQ domains of IQD14 that are distinct from those observed with CaM. For example, in the case of CML13, there is a decrease in interaction affinity in the presence of Ca^2+^ (Table 2). In general, the interactions of CaM, CML13, and CML14 with IQ-domain peptides from IQD14 were driven by favourable enthalpic contributions with unfavourable changes in entropy (Supplementary Information: Tables S4, S5). However, in several cases binding events were driven by both favourable entropy and enthalpy. For example, in the lower-affinity binding event for CML13’s association with IQD14-IQ1+2 in the presence of EGTA (*Ka2*, Supplementary Information: Table S4), and in the higher-affinity binding event for CaM’s association in EGTA with IQD14-IQ2 (*Ka1*, Supplementary Information: Table S4), both favourable enthalpic and entropic contributions were observed (Supplementary Information: Tables S4, S5).

Given the complexity of the interactions suggested by ITC analyses, we sought to further explore the stoichiometry of CaM/CML binding to IQD14-IQ1+2 via sedimentation profiles using analytical ultracentrifugation (AUC). The binding of either CaM, CML13, or CML14 to IQD14-IQ1+2 were observed at a 1:1 stoichiometry based upon the computed molecular weight for the individual proteins and the observed complexes (Table 3). The AUC data thus support the predicted 1:1 stoichiometry from the ITC analysis and indicate that, even with two IQ domains represented in the IQD14-IQ1+2 peptide, only a single CaM or CML protein was associating under our experimental *in vitro* conditions.

**Table 3.** Analytical ultracentrifugation analysis of CaM, CML13, and CML14 interactions.

| **Protein analysed<sup>1</sup>** | **Apparent molecular weight<sup>2</sup>** | **Frictional ratio<sup>3</sup>** |
| --- | --- | --- |
| IQD14 IQ1+2 | 5.7 kDa | 1.20 |
| CaM | 17.0 kDa | 1.32 |
| CaM + IQD14-IQ1+2 | 22.2 kDa | 1.28 |
| CML13 | 19.6 kDa | 1.28 |
| CML13 + IQD14-IQ1+2 | 25.1 kDa | 1.13 |
| CML14 | 22.2 kDa | 1.32 |
| CML14 + IQD14-IQ1+2 | 27.6 kDa | 1.30 |

<sup>1</sup>Listed are the proteins and/or synthetic peptides used in each sedimentation velocity experiment.
<sup>2</sup>Apparent molecular weight as determined by SEDFIT v16.1.
<sup>3</sup>Frictional radius associated with the reported molecular weight in SEDFIT.

### Circular dichroism spectroscopy (CDS) of IQD14 binding to CML13, CML14, and CaM

CDS was used to further examine the structural properties of CaM/CML/IQ interaction. Far-UV CD spectra of CML13 and CML14 were nearly unchanged in the presence or absence of Ca^2+^, whereas CaM exhibited notable secondary structural changes (Figure 10A). Not surprisingly, the IQD14-IQ1+2 peptide alone appeared 92% unstructured by far-UV CD. We compared the actual spectrum for the complex of CaM, CML13 or CML14 with the IQD14-IQ1+2 peptide to the theoretical spectrum derived from the summation of the separate CaM/CML and peptide spectra. We observed a gain in helical content as shown by increased signals at the 3 peaks indicative of alpha helical content (190, 208, 222 nm). The differential spectrum, derived by subtracting the theoretical from the actual spectrum, highlights this observation (Figure 10A). This gain of structure by the peptide, which indicates interaction with CaM or the CMLs, did not require Ca^2+^. These *in vitro* data provide further support for CaM/CML-IQ domain interaction and corroborate the Ca^2+^-independent binding we observed using ITC and spot-blot overlay assays.

**Figure 10.**
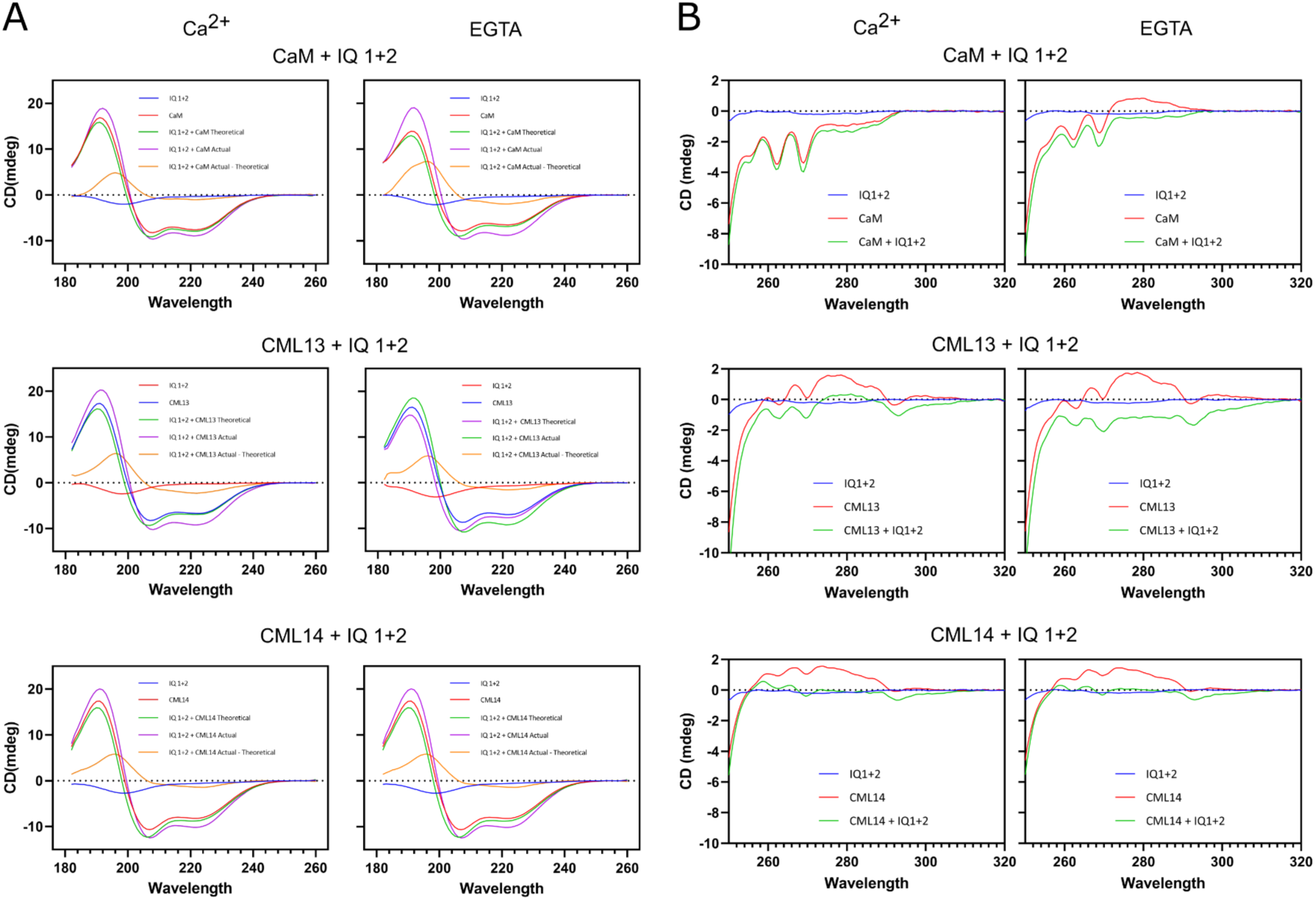
Analysis of structural changes in CaM, CML13, CML14 during interaction with an IQD14 synthetic peptide. Far-UV CD spectroscopy (**A**) was used to explore secondary structure, and near-UV spectroscopy (**B**) was used to examine tertiary structure of the synthetic peptide IQD14-IQ1+2 individually or in complex with recombinant CaM, CML13, or CML14 in 2 mM CaCl2 or 2 mM EGTA as indicated. See Fig 9A for the peptide sequence and corresponding position in IQD14. Theoretical interaction spectra were calculated by addition of individual spectra therein. Protein concentrations were 58 μM, 59 μM, 48 and 57 μM for CaM, CML13, CML14, IQ 1+2 peptide, respectively (far-UV), and 60.75 μM, 55.3 μM, 57 μM and 53.5 μM for Cam, CML13, CML14, and IQD14 1+2, respectively (near-UV). Each plot is an average of 10 scans measured in 1 nm units from 183-260 nm (far-UV) with a 0.1 cm pathlength or 0.5 nm units from 240-320 nm (near-UV) with a 1 cm pathlength.

To further investigate changes in tertiary structure during CaM/CML-IQ domain binding in the presence or absence of Ca^2+^, we utilized near-UV CDS (Figure 10B). Although slight differences exist between the near-UV spectra of CML13 and CML14 in complex with IQD14-IQ1+2, their respective spectra in the presence of Ca^2+^ vs EGTA are nearly identical, confirming a lack of Ca^2+^ dependence in their interaction with the peptide. Under either Ca^2+^ or EGTA conditions, we observed a shift in the 275-282 nm region and around 290 nm, suggesting spectral contribution from the single Tyr and single Trp residues, respectively, present in the C-lobe of both CMLs. In contrast, CaM/IQD14-IQ1+2 complexes displayed clear differences in near-UV spectra in the presence of EGTA vs Ca^2+^, indicating distinct binding properties in response to Ca^2+^. A tertiary change in CaM was observed for CaM/peptide interaction but only under EGTA (apo) conditions. In contrast, the tertiary change in Ca^2+^-bound CaM was minimal upon peptide binding. This may suggest that when CaM in is the holo-form, the structural changes that occur in the C-lobe (see Figure 3) precede peptide binding, whereas the apo-form undergoes more pronounced tertiary changes in response to peptide binding. Taken together, CDS analyses support the interaction data from ITC and suggest that in comparison to CaM/peptide interaction, any conformational changes in tertiary structure by CML13 or CML14 that occur upon binding to the IQ1+2 peptide are much less pronounced and below detection by near-UV CDS.

## DISCUSSION

CMLs are the largest family of Ca^2+^ sensors in plants, yet only a few putative targets of CMLs have been described (Dobney et al., 2009; La Verde et al., 2018; Pedretti et al., 2020). Plant CAMTAs, IQDs, and myosins are known targets of CaM (Bouche et al., 2002; Poovaiah et al., 2013; Haraguchi et al., 2014), and our data indicate CML13 and CML14 are also candidate partners of these proteins. In comparison to CaM, CML13 and CML14 displayed a binding preference for tandem IQ domains. Nevertheless, CML13 and CML14 were capable of interacting with some of the single IQ domains of IQD14 *in vitro*, albeit with lower affinity compared to tandem IQ domains (Figures 5, 7, Table 2). That we observed almost no growth when using the Y2H system, but a weak signal in the split-luciferase assays for CML13, CML14 when using the single-IQ domain from CNGC20, may be due to greater sensitivity in the latter method. Thus, although CML13 and CML14 are capable of binding to at least some single IQ domains, and possibly other types of CaM-binding domains, they may show a preference for targets with tandem IQs and further study into their specificity is merited. Among the 12 different CaM/CMLs tested, only CaM, CML13, and CML14 provided a strong interaction signal with IQD14 in the *in planta* split-luciferase system (Figure 5B). The structural properties of CaM and CML13, and CML14 and/or cell conditions underlying this specificity remain unknown. However, as we did observe weak but detectable signals relative to CaM, CML13, CML14 when testing various CMLs *in planta* (Fig 5B), and when testing CML42 binding to IQD14 *in vitro* (Supplementary Information: Figure S6), it remains possible that, aside from CML13 and CML14, other family members may associate with IQ-domain proteins under specific *in vivo* conditions. That said, the strong affinities (nM range) that we observed for CML13 and CML14 for IQ domains, and the fact that CML13 and CML14 are typically the most abundant CMLs in proteomic databases (Supplementary Information: Table S1), strongly suggests that CML13 and/or CML14 would likely outcompete other CMLs for binding to CAMTAs, IQDs, and myosins.

Additional questions remain on how Ca^2+^ might impact CaM, CML13, and CML14 association with IQ-target proteins *in vivo*. While both the Y2H and split-luciferase systems offer the advantages of cell-based assays, manipulating internal Ca^2+^ levels over the duration of growth is not practical, and thus it remains unclear how Ca^2+^ signals may contribute to *in vivo* CaM/CML interaction with IQ-targets. Our *in vitro* ITC and overlay assays suggest while Ca^2+^ is not essential for interaction, it likely alters binding affinities and/or specificity and thus further study into CML/IQ-target specificity and the impact of Ca^2+^ is merited.

Several studies have proposed that IQDs may be targets of some CMLs (Yong-Yeol Ahn et al., 2011; Bürstenbinder et al., 2013; Kölling et al., 2019), and Wendrich et al. (2018) observed CaM, CML13, and CML14 among the various proteins that co-immunoprecipitated with several Arabidopsis IQD proteins from crude extracts, supporting our hypothesis that IQDs are *in vivo* targets of these CMLs. It is also worth noting that CaM-related MLCs that bind to IQ-domains have been characterized in other organisms (Crawley et al., 2006; Crawley et al., 2011; Langalaan et al., 2016). Furthermore, CML24 was reported as a putative interactor of myosin ATM1 (Abu-Abied et al., 2006), and Haraguchi et al., (2018) speculated that, in addition to CaM, CaM-related proteins may function as MLCs. In the case of CAMTAs, there is evidence that they interact with CaM *in vivo*, and that their IQ domains play key functional roles (Iqbal et al., 2020) and a recent report showed interaction between CML24 and the C-terminal region of CAMTA2 during aluminum stress in Arabidopsis (Zhu et al., 2022).

Our data indicate that CML13 and CML14 display minimal conformational change upon Ca^2+^ binding (Figure 3), consistent with previous observations on CML14 (Vallone et al., 2016). Both CML13, and 14 have several unusual residues in (degenerate) EF hand-3 and -4 in comparison to CaM and other CMLs, and EF-hand 2 is highly degenerate, with gaps within the consensus sequence (Figure 2). Although the consensus sequence of EF hand-1 in CML13 and CML14 are identical, two residues immediately downstream are not conserved (ES vs QA), and these changes may explain the differences between CML13 and CML14 in Ca^2+^ sensitivity we observed using ITC. Previous studies indicate that the Ca^2+^ binding affinity of CaM or some CMLs can be altered in the presence of a target peptide (Gifford et al., 2007; Pedretti et al., 2020). Thus, we tested whether IQD14 peptide binding altered the global Ca^2+^ affinity of CML13 *in vitro* but we did observe a major change (Supplementary information: Fig S7). Although roles for CML13 and CML14 as direct Ca^2+^ sensors seem unlikely, it is possible that they may complement the function of CaM by interacting with IQ-domain targets under certain conditions and could thus be considered components of Ca^2+^-based signal transduction in a broader sense. For example, CML13 and CML14 may compete with CaM for binding to some IQ domains. The quantity of “free” CaM (i.e. not bound to a target) in the cytosol will vary with cell conditions, but will drop upon Ca^2+^ influx when CaM is occupied with an array of targets (Black et al., 2004). A drop in free [CaM] that causes dissociation of IQ-bound CaM from some targets, or even specific IQ domains, could facilitate CML13 and/or CML14 preserving the role of an IQ-bound regulator while also increasing the quantity of free CaM available to the cell. Such a model remains speculative but is consistent with reports indicating free [CaM] may be limiting under certain conditions (Persechini & Cronk, 1999; Black et al., 2004). Moreover, the broad expression pattern of CML13 and CML14 is reminiscent of CaM, whereas many CMLs are often tissue-specific or responsive to external stimuli (McCormack & Braam, 2005; Dobney et al., 2009; Ogunrinde et al., 2017). It is also worth noting that CML13 and CML14 are typically the most abundant CMLs described in public Arabidopsis proteome databases (Baerenfaller et al., 2011), which may position them as available partners for IQ targets that also bind CaM. This model would predict a reversible state, dependent upon cell conditions, free concentrations of CaM/CML, and other factors. Alternatively, some IQ domains may be CML13- and/or CML14-specific, and thus not compete with CaM, as is the case with several CaM-related, Ca^2+^-insensitive MLCs in other organisms (De La Roche et al., 2003; Heissler & Sellers, 2014). Regardless, given the high level of phylogenetic conservation (Zhu et al., 2015), it seems likely that CML13 and CML14 orthologs target similar IQ-domain proteins across plant taxa.

It is also possible that CML13 and CML14 may play indirect roles in Ca^2+^ signalling by modulating binding partner activity in conjunction with CaM or other CMLs. IQD proteins function as scaffolds at the cytoskeleton, affecting cell morphology and development (Burstenbinder et al., 2017). For example, IQD5 participates in CaM-dependent protein-protein interactions with kinesin light-chain related protein, with CaM serving as a molecular scaffold (Zhu et al., 2015; Mitra et al., 2019). CMLs might therefore affect the recruitment of IQD binding partners, reminiscent of the proposed role for CML42 in modulating KCBP (kinesin-like CaM binding protein) binding to microtubules through interactions with KIC (KCBP-interacting Ca^2+^ binding protein; Kölling et al., 2019).

The localization of CML13 and CML14 to the cytosol and nucleus (Figure 6A) is consistent with that of their putative targets; CAMTAs (nuclear; Kidokoro et al., 2017), IQDs (microtubules and/or nuclear; Burstenbinder et al., 2017), and myosins (actin cytoskeleton; (Golomb et al., 2008; Peremyslov et al., 2012; Haraguchi et al., 2014). Moreover, CML13, CML14, and CaM relocalized to microtubules when coexpressed with IQD14 in tobacco cells, providing additional support that these interactions can occur *in vivo* (Figure 6B).

Although our *in planta* and *in vitro* binding data indicate specific interaction of CML13 and CML14 with representative members of the IQD, myosin, and CAMTA families, the situation *in vivo* may be quite complex. The estimated binding affinities of CML13, CML14, and CaM with a synthetic peptide from IQD14 with two IQ domains are comparable (nM range) to those observed for other CaM or CaM-related proteins (DeFalco et al., 2016) and are thus likely within the physiological range. However, the binding kinetics within a cell will be influenced by a range of factors that are not readily tested in the controlled *in vitro* environment. For example, microdomains of spatially-localized Ca^2+^, CaM, or CMLs, could impact the specificity and affinity of IQ partners. Similarly, post-translational modifications of either CML or IQ-domain partner could add another layer of complexity. Indeed, although CML13 and CML14 are 95% identical, of the seven residue differences, several have been identified as differentially phosphorylated *in vivo* based on public phosphoproteome data (Willems et al., 2019). Moreover, it is possible that different combinations of CaM and/or CMLs bind to distinct IQ domains within a single protein target *in vivo*. With 6, 17, and 33 members, respectively, among CAMTAs, myosins, and IQD proteins in Arabidopsis, and the number of paired-IQ domains also ranging within family members, the hypothetical complexity of CaM/CML interaction with any given IQ-target is substantial.

In general, our binding analyses using CML13 and CML14 support previous studies using CaM or CaM-related proteins that showed markedly different affinities for pairs vs single IQ domains (Langelaan et al., 2016; Powell et al., 2017). These data also help to explain the apparent contradiction of IQD14-IQ2+3 interacting with CaM/CMLs in the Y2H system whereas single IQ2 and IQ3 domains do not, an observation that likely also reflects the greater sensitivity of ITC for protein-protein interaction in comparison to Y2H studies. In general, the underlying mechanisms of CaM, CML13, and CML14 binding are difficult to unravel without solved structures given that interactions with multiple IQ domains are known to be complex (Li & Sacks, 2003; Andrews et al., 2021). The ITC data for CML13 and CML14 appear to be inherently simpler than those for CaM, and this may reflect the fact that CaM has four functional EF-hands. The IQD14-IQ1+2 binding curves display an asymmetry which suggests competition between two binding conformations, and not simply two binding events, when two IQ domains are present (Figure 8), a phenomenon previously described (Brautigam et al., 2016). The thermodynamic properties of the interaction of CaM and CML with the IQD14 peptides are reminiscent of previous reports showing favourable enthalpic and unfavourable entropic contributions to CaM binding to some targets (Dunlap et al., 2013; Taiakina et al., 2013). In general, CaM-binding domains that undergo changes from unstructured to a structured helical conformation upon CaM binding are more likely to be driven by favourable enthalpy (Dunlap et al., 2013). Our CDS data, indicating that the IQD14 peptides are unstructured in the absence of CaM or CML, is consistent with this interpretation. However, it should be noted out that although the synthetic peptides we used possessed peripheral regions outside of the IQ domains, it remains possible that additional residues, not included in our analysis, may impact the thermodynamics and kinetics of CaM/CML binding to target IQ domains. The poor solubility of recombinant proteins with multiple IQ domains, regardless of expression system, and the size limitations of synthetic peptides, create substantial challenges for *in vitro* binding analysis using IQ targets (Homma et al., 2000).

It is worth noting that within the IQ67 domain of IQDs there appears to also be overlapping 1-5-10, and 1-(5)-8-14 motifs (corresponding to the positions of hydrophobic residues), which are considered typical CaM-binding motifs (Abel et al., 2005; Andrews et al., 2021). One possible binding mechanism is described by the Ca^2+^ “switch” model of CaM in association with an IQ domain from mouse myosin Va (Shen et al., 2016). In this model, the C-lobe of CaM binds in a semi-open state through hydrophobic and electrostatic interactions and the N-terminal lobe of CaM is in the closed state and is weakly bound. In the presence of Ca^2+^, the N-lobe of CaM rotates to the opposite side of the myosin IQ α-helix as a result of interactions in the linker region of CaM. In our analysis, we may be observing two binding events, one from each lobe of CaM, rather than binding and rearrangement, and while this is speculative, similar observations have been reported (Li et al., 2017). Indeed, we tested the interaction of the individual N- and C-lobes for each of CaM, CML13, and CML14 with the IQ-region of IQD14 using the split-luciferase system (Supplementary Information: Figure S8). Among the N-lobes, only CML13 showed a strong interaction signal with IQD14, whereas the C-lobes for all three bait proteins gave strong binding signals. Nevertheless, we cannot exclude the possibility that the *in vivo* binding mechanism may involve both lobes of these proteins under cellular conditions, and our data do not allow us to confidently state whether the multiple binding events that we observed in ITC analysis correspond to sequential lobe binding or complex rearrangement.

Although the *in vivo* functions of CML13 and CML14 remain unclear, and may vary depending upon their targets, studies of CaM and CaM-related proteins in association with IQ domains of animal myosins may offer insight. CaMs and CaM-related proteins function as MLCs and play both structural and regulatory roles by binding and stabilizing the IQ-domains of the neck region, and if MLCs become unbound, myosin activity is impaired or arrested (Heissler & Sellers, 2014). There are 17 myosins in Arabidopsis and while CaM functions as an MLC for certain isoforms, some myosins do not show *in vitro* motility when using CaM alone as the MLC (Haraguchi et al., 2018). We speculate that CML13 and CML14 may function as novel plant MLCs, reminiscent of the CaM-related MLCs observed in other organisms (Heisler & Sellers; 2014). Interestingly, the moonlighting of MLCs as interactors with cytoskeletal scaffolds is a known phenomenon, as the human IQ-motif containing GTPase activating proteins (IQGAPs) and yeast IQGAP-like proteins perform similar functions to IQDs and interact with the essential light chains Mlc1sa and Mlc1p, respectively (Boyne et al., 2000; Pathmanathan et al., 2008; Atcheson et al., 2011). Although the *in vivo* function of the IQGAP Mlc1sa interaction is still unknown, the Mlc1p interaction with both IQGAP-like and myosin II proteins at the contractile ring are crucial for proper cytokinesis (Boyne et al., 2000). Since both IQDs and myosin VIIIs have recently been implicated in preprophase band formation and cellular division (Wu & Bezanilla, 2014; Kumari et al., 2021), it is possible that CML13 and CML14 in plants play a similar role to that reported previously for Mlc1p.

In the case of CAMTAs, the CaM/CML interactions are likely complex given that CAMTAs possess both tandem IQ domains and a separate C-terminal, CaM-binding domain (Bouché et al., 2002). CAMTAs are key regulators of transcription in plants, linking Ca^2+^ signals to changes in gene expression (Liu et al., 2015). Mutations in the CaM-binding or IQ domains of CAMTA3 have direct implications on mutant phenotypes and CAMTA3-regulated gene transcription (Kim et al., 2017), and a recent study indicates that CML24 binding to CAMTA2 promotes transcription of *ALMT1* in response to aluminium stress (Zhu et al., 2022). These results suggest that both CaM and CMLs could be acting in concert to interpret Ca^2+^-signals and alter CAMTA-dependent gene transcription via the C-terminal IQ and CaM-binding domains in response to stimuli. With various myosins, CAMTAs, and IQD members as potential CML13,14 targets, the functional reach of these CMLs is hypothetically quite broad, from cytoskeletal organization to gene expression, and will require a range of strategies and additional studies in the future to uncover.

## EXPERIMENTAL PROCEDURES

### Plant Growth Conditions

*Arabidopsis thaliana* (Arabidopsis) Col-0 or *Nicotiana benthamiana* seeds were sown into Sunshine mix #1 (Sun Gro Horticulture Canada Ltd.) and stratified at 4°C for 3-5 days in the dark before transfer to a growth chamber (Conviron MTR30) under long (18 h) or short (12 h) photoperiod conditions, for Arabidopsis and *N. benthamiana*, respectively, at 22°C, ∼150 *μmol* m^-2^ s^-1^. For growth on sterile agar plates, Arabidopsis seeds were surface sterilized using the ethanol/bleach method (Weigel et al., 2002) and spread onto 0.5x MS basal salts with macronutrients and micronutrients (Murashige & Skoog, 1962; Caisson Laboratories Inc.), 0.8% (w/v) agar plates followed by stratification and growth in a chamber as described above.

### Gene annotations

Arabidopsis locus codes for genes used in this study are: At4g03290 (*CML6*), At4g14640 (*CML8*), At1g12310 (*CML13*), At1g62820 (*CML14*), At1g18530 (*CML15*), At4g37010 (*CML19*), At1g66400 (*CML23*). At5g37770 (*CML24*), At2g41410 (*CML35*), At1g76650 (*CML38*). At4g20780 (*CML42*), At2g43680 (*IQD14*), At3g17700 (*CNGC20*), At5G17330 (*GAD1*), At1G67310 (*CAMTA4*), At4G27370 (*myosin VIII-B*), At4G33050 (*IQM1*). For all experiments involving CaM, the cDNA encoding petunia CaM81 (GenBank, M80836.1) was used (Fromm and Chua, 1992). CaM81 is an evolutionarily-conserved isoform, hereafter termed CaM, that is 100% identical at the amino-acid sequence level to Arabidopsis AtCaM7 (At3g43810) and differs by a single residue to conserved AtCaMs 1,2,3,5,6 (McCormack & Braam, 2003).

### Histochemical GUS Assays

Histochemical staining to detect GUS activity in the *CML13*pro::GUS and *CML14*pro::GUS transgenic plants was performed using 5-bromo-4-chloro3-indolyl-β-D-glucuronide (X-gluc, BioShop) as a substrate (Jefferson et al., 1987; Vanderbeld and Snedden, 2007). Leaf tissue was harvested and immediately fixed in ice-cold 90% acetone for 30 min. The tissue samples were then washed with distilled water and incubated in GUS staining solution (100 mM Na2HPO4-NaH2PO4 buffer pH 7.0, 0.5 mM EDTA, 0.1% (v/v) Triton X-100) for 2 h. X-gluc was added to the GUS staining solution for a final concentration of 0.5 mM. The tissue was washed several times with 70% ethanol over a period of 24 h to clarify the tissue. Stained tissue samples were examined and digitally photographed using a ZEISS SteREO Discovery V.12 dissecting scope attached with an EMS-1 controller and a Nikon Coolpix P7800 camera.

### Plasmid Vector Construction

PCR amplification was done using Q5 DNA polymerase and manufacturer guidelines (New England Biolabs Ltd.) and all plasmid constructs were generated by standard molecular cloning techniques and confirmed by Sanger DNA sequencing (TGCA Sequencing Centre, The Hospital for Sick Children, Toronto). Oligonucleotide primer sequences and information are listed in Supplemental Table S3. GUS promoter:reporter constructs pBI101-*CML13*pGUS and pCambia1305.1-*CML14*pGUS were made by PCR amplification of the 1.5 kb and 1.01 kb regions upstream of the first ATG codon of *CML13* and *CML14*, respectively.

For subcellular localization experiments, full-length open reading frames encoding CaM, CML13 and CML14 were subcloned into pRLT2ΔNS/mGFPx2 using SalI and EcoRI (CaM) or HindIII and EcoRI (CML13 and CML14) restriction sites. The full-length open reading frame of IQD14 was subcloned into pRTL2/mcherry using the KpnI and NheI (using XbaI compatible ends) restriction sites. The construction of pRTL2/NLS-RFP, encoding three tandem copies of the nuclear localization signal (NLS) fused to red fluorescent protein (RFP), has been described previously (Dhanoa et al. 2010). For the split-luciferase protein-interaction assays, full open-reading frames for all CaM or CML bait proteins were subcloned in-frame as N-terminal fusions with the C-lobe of firefly luciferase using the pCambia1300-CLuc vector (Chen et al., 2008). All other putative interactor prey proteins were cloned in-frame as C-terminal fusion proteins with the N-lobe of firefly luciferase using the pCambia1300-NLuc vector. See Figure Legends for the specific regions of bait or prey proteins tested for interaction.

### Yeast 2-hybrid (Y2H) cDNA Library Screen

A Y2H library screen was performed as outlined in the Matchmaker Gold “Mate and Plate” Yeast Two-Hybrid System (Takara Bio Inc.) using pGBKT7-CML13 as the bait in yeast strain AH109. Serial dilutions were plated onto single-dropout (SD) auxotrophic-selection media lacking either –Leu, -Trp, or -Leu/Trp (SD-L,-T,-LT) to check the transformation efficiency and 200 µL was plated onto 55 x 150 mm SD-Leu/Trp/His/Ade (SD-LTHA) plates and allowed to grow at 30°C for 3-5 days. Approximately 100 independent yeast colonies were cultured for analysis of the respective prey plasmids. Prey plasmids were isolated using the Presto™ Mini Plasmid Kit (Geneaid Biotech Ltd.) and putative CML13-interactors which were found not to self-activate the reporter genes by pairwise transformation were identified by Sanger DNA sequencing. For pairwise interaction Y2H tests pGBKT7-CML13, pGBKT7-CML14 and pGBKT9-CaM were used as bait vectors and chemically-competent yeast (AH109) cells were prepared and transformed using standard LiAc/PEG mediated protocols (Schiestl and Gietz, 1989), an aliquot (100 µL) of transformed cells was then plated onto SD media containing the appropriate dropout supplement (Takara Bio USA, Inc.), and grown at 30°C for up to 7 days.

### Bioinformatic Analysis

Gene expression patterns for *CMLs* of interest were obtained from public database resources using Arabidopsis eFP Browser, Bio Analytic Resource (BAR, University of Toronto). Raw *CML* expression values were normalized to actin and analysed. Publicly available proteomic data were obtained from the PaxDb database (Wang et al. 2015). Multiple sequence alignments were performed using ClustalΩ (Sievers et al., 2011) and were edited for images using BIOedit software (Hall T.A., 1999).

### Split-Luciferase Assay

The split-luciferase assays were performed essentially as described (Chen et al., 2008). Four-week-old *N. benthamiana* leaves were infiltrated by *Agrobacterium tumefaciens* (Agrobacterium) strain GV3101 containing the plasmids of interest. Whole leaves were removed after 4 d incubation and sprayed with a 1 mM D-Luciferin (GoldBio) and 0.01% Silwet L-77 (Lehle Seeds Inc) solution followed by incubation in the dark for 10 min and imaging using a ChemiDoc™ Touch System (Bio-Rad). Alternatively, leaf discs were taken after 4 d, incubated with 100 μL water containing 1 mM D-luciferin in a 96-well plate for 10-15 min, and luminescence was captured with the SpectraMax Paradigm multimode detection platform (detecting all wavelengths, 10 s integration time; Molecular Devices Inc.) with LUM module. Expression of CaM/CML C-Luciferase fusion bait proteins was verified by immunoblotting (Santa-Cruz Biotech).

### Recombinant Protein Expression, Processing and Quantitation

Recombinant proteins were expressed in *E. coli* BL21 strain pLysS for CaM or CPRIL (Novagen) for all CMLs. Bacterial extracts were prepared using a French Press Cell (Glen Mills Inc), and purified as described (Dobney et al., 2009). Soluble bacterial lysate was collected and flash frozen in liquid N2 or used immediately for downstream processing. For processing of insoluble proteins, insoluble bacterial lysate was separated by centrifugation and the pellets were solubilized in 8 M urea. His8-tagged soluble or insoluble CML recombinant proteins were purified at room temperature using Ni-NTA resin (His60, Takara Bio Inc.) using the manufacturer’s protocol. Recombinant CaM was purified using phenyl-sepharose resin (*Phenyl Sepharose*^®^ 6 Fast Flow, GE Healthcare Inc.) as described (Bender et al., 2013). Prior to downstream analysis, purified proteins were concentrated using AMICON centrifugal filters (MWCO 10-30 kDa, MilliporeSigma Inc.) and dialysed into appropriate buffers for downstream analysis. Proteins were quantified using either the Bradford assay (BioShop Canada Inc.), Bicinchoninic acid (BCA) assay (Bio Basic Inc.), or, for all biophysical analyses, by commercial Amino Acid Analysis (AAA; SPARC labs, The Hospital for Sick Children, Toronto).

### *In vitro* Protein-Interaction Overlay Assays

CaM/CML overlay assays were performed as previously described (DeFalco et al., 2010) with minor alterations. CaM or CMLs were either labelled with biotin-NHS Ester (ThermoFisher) or with fluorescent probe 680RD (LI-COR) as per the respective manufacturer’s instructions. Triplicate aliquots (200 ng) of putative target proteins, or BSA controls, were spotted onto nitrocellulose blots, allowed to dry, then blocked in 5% casein and incubated for 1 h at room temperature in TBS-T with 2 mM CaCl2 or 5 mM EGTA and 200 nM of the respective biotinylated or fluorescently-tagged CaM/CMLs. Blots were then washed in the same TBS-T buffer without CaM/CMLs and directly detected using a LI-COR Odyessy XF imager or, for biotin-labelled probes, incubated 1 h in Streptavidin-HRP (Sigma) as per manufacturer’s instructions, followed by ECL chemiluminescence (Bio-Rad) detection and imaged using a ChemiDoc™ Touch Imaging System (Bio-Rad).

### SDS-PAGE and Western Blotting

SDS-PAGE, protein visualization using Coomassie R-250, and immunoblotting was performed as described using standard protocols (Bender et al., 2013). For immunoblotting, proteins were transferred to nitrocellulose membranes and blocked using 5% (w/v) BSA in TBS-T. Membranes were incubated with primary antibody for one hour (α-CLuc; Santa-Cruz Biotech) at room temperature or overnight at 4°C (α-His; GenScript), followed by washes with TBS-T, incubation with horseradish-peroxidase-conjugated secondary antibody (Sigma) for one hour at room temperature, and detection using Bio-Rad’s Clarity™ Western Blotting Substrates and a ChemiDoc™ Touch Imaging System (Bio-Rad).

### Synthetic Peptides

Synthetic peptides at >85% purity for IQD14-IQ1, residues 322-346 (Nt-QHASATKIQGAFRGYMARKSFRALK-Ct), -IQ2, residues 344-367 (Nt-ALKGLVRLQGVVRGYSVKRQTINA-Ct), and -IQ1+2, residues 322-367 (Nt-QHASATKIQGAFRGYMARKSFRALKGLVRLQGVVRGYSVKRQTINA-Ct), all with N-terminal acetylation and C-terminal amidation, were purchased commercially (Bio Basic Inc), solubilized in buffers suitable for analyses (see figure legends) and used fresh or flash frozen in liquid nitrogen and stored at –80°C. A similar nomenclature applied to all peptides in the study. For example, the annotation “IQD14-1+2” denotes a peptide with the first and second IQ domains of IQD14, positioned relative to the N-terminus, whereas “IQD14-IQ2” refers to the second IQ domain alone. Peptide concentrations for analyses were determined by AAA (SPARK labs, The Hospital for Sick Children, Toronto).

### Isothermal Titration Calorimetry (ITC)

ITC was performed at the Queen’s University Protein Function Discovery (PFD) facility as described (DeFalco et al., 2010) using an AVP-ITC colorimeter (Microcal, LLC) at 30°C with ITC buffer (10 mM Tris-HCl, pH 7.5, containing 2 mM CaCl2 or 2 mM EGTA).

Synthetic peptides for binding assays were loaded in the ITC cell at 15, 30, or 45 µM under conditions described in the figure legends. Forty, 7.5 µL injections of CaM, CML13, or CML14 (see figure legends for sample concentrations) were performed in 300-second intervals. Samples were degassed and brought to 30°C prior to loading. Data was fit to a one- or two-site model using Origin 7.0 software (OriginLab Corp.).

### Analytical Ultracentrifugation (AUC)

AUC analysis was performed at the Queen’s University PFD facility. AUC was used to estimate the stoichiometry of the CaM/CML IQD14-IQ1+2 complex. Protein samples (CaM, CML13, CML14, and IQD14 1+2 peptide at concentrations of 60.75 µM, 55.3 µM, 57 µM, and 53.5 µM respectively) were dialyzed into 10 mM Tris-Cl, (pH 7.5) and supplemented with CaCl2 or EGTA to a final concentration of 2 mM. AUC was performed at 20°C in a Beckman Optima XL-I analytical ultracentrifuge with 12 mm, 2-sector Epon-charcoal cells, and an An-60 Ti rotor. Sedimentation behavior resulting from centrifugation at 50 000 RPM was observed using the optical absorbance and interference optics of each sample at a wavelength of 280 nm. The initial 200 scans obtained from each cell were fitted according to the continuous c(S) Lamm equation model in the SEDFIT software package (version 9.4) in order to obtain the relative molecular weights.

### Circular Dichroism (CD) Spectrometry

Near- and far-UV CD spectroscopy, used to evaluate structural features of CaM, CML13, and CML14 with or without IQD14-IQ1+2 binding peptide, was performed using a Chirascan V100 CD spectrophotometer with adaptive scanning and a cylindrical quartz cuvette with a pathlength of 0.1 mm (far-UV) or 1 cm (near-UV). Protein samples (see figure legend for concentrations) were prepared by dialysis overnight into 10 mM Tris-Cl, (pH 7.5) and supplemented with CaCl2 or EGTA to a final concentration of 2 mM. Spectra from 10 scans were averaged per assay. The differential absorption of left and right circularly-polarized light is presented as raw absorbance or molar ellipticity (θ) in degrees. Average corrected dichroism spectra for CaM, CML13 and CML14 alone were deconvoluted using both the OLIS GlobalWorks and CDNN deconvolution software to reveal percentage of secondary structure.

### Ca^2+^ titration binding assay with chromophoric chelator

Ca^2+^ binding to CML13 was assessed using titration analysis as previously described (Pedretti et al., 2020), based on the competition for Ca^2+^ between the protein and the chromophoric chelator 5,5′-Br_2_-BAPTA [5,5’-dibromobis-(o-aminophenoxy)ethane N,N,N’,N’-tetra-acetic acid] (Sigma). All solutions were decalcified using Chelex-100 resin (Sigma) as per to manufacturer’s instructions. Ca^2+^ titrations, from 0-50 μM, were performed by combining a similar concentration of CML13 and chelator in 5 mM Tris-HCl, 150 mM KCl, pH 7.5. Chelator concentration was 28 μM and the initial Ca^2+^ concentration was 2.8 μM. CML13 protein concentration (20-24 μM) was measured by the Bradford assay. The absorbance at 263 nm (λ_max_ for the Ca^2+^ free chelator) was measured as 5 μL aliquots of 1 mM CaCl_2_ were added until no significant changes in absorbance was detected. Ca^2+^ titrations were performed with excess IQD14-IQ1 peptide using a protein:peptide ratio of 1:2. CaLigator software (Andre et al., 2002) was used for data analysis, using a model with a chelator of known Ca^2+^ affinity ([K_d_ (5,5′-Br_2_-BAPTA) of 2.3 μM in 0.15 M KCl] and 2 macroscopic Ca^2+^ binding constants for the protein (log K_A_). The apparent affinity value (K_d_) was determined from the average of the logarithms of the four macroscopic binding constants (K_d_ = 10 ^(logK1+logK2)/2)^) as described (Andre et al., 2002).

### BY-2 cell transformation and microscopy

The preparation and use of tobacco (*N. tabacum*) BY-2 suspension-cultured cells and biolistic bombardment using a PDS-1000/He biolistic particle delivery system (Bio-Rad Laboratories) were performed as previously described (Banjoko and Trelease, 1995; Teresinski et al., 2019). Bombardments included either 2 µg of pRTL2/NLS-RFP, serving as a nuclear marker protein (Dhanoa et al., 2010) or 4 µg of pRTL2/IQD14-mCherry and 4 µg of either pRTL2/GFPx2, pRTL2/CaM-GFPx2, pRTL2/CML13-GFPx2, or pRTL2/CML14-GFPx2. Following bombardment, cells were incubated at 25 °C in the dark for ∼5 h to allow for gene expression and subcellular sorting of the introduced fusion proteins. Cells were then formaldehyde-fixed (Banjoko and Trelease, 1995) and images of cells were acquired using an Axioskop 2 MOT epifluorescence microscope (Carl Zeiss Inc.) equipped with a Zeiss 63X Plan Apochromet oil-immersion objective and a Retina 1300 charged-couple device camera (Qimaging) using Open Lab software package (version 5.0) (Improvision Inc.). Images were deconvoluted using the Diffraction PSF 3D plugin (https://www.optinav.info/Diffraction-PSF-3D.htm) and the Parallel Spectral Deconvolution 2D 1.9 plugin (https://imagej.net/Parallel_Spectral_Deconvolution) for ImageJ 1.53e (Schneider et al., 2012). All fluorescence images shown are representative of >15 co-transformed cells from at least two separate bombardments. Figure compositions were generated using Adobe Photoshop CS (Adobe Systems).

## Data Availability Statement

All relevant data can be found within the manuscript and its supporting materials.

## Supporting information

Combined Supp Figs and Tables

## Acknowledgements

The authors would like to thank Dr. Jacqueline Monaghan (Queen’s University) for helpful comments on the manuscript.

## Short Legends for Supplemental Figures

Supplemental Figure S1. Transcriptomic databases indicate CML13 and CML14 have broad expression patterns.

Supplemental Figure S2. Representative, whole-leaf images of split-luciferase CML/IQD14 protein-protein interaction assay.

Supplemental Figure S3. Immunoblot showing expression of CML-bait proteins in *N. benthamiana* split-luciferase protein interaction assays.

Supplemental Figure S4. Y2H delineation of interaction region between Arabidopsis CML13/14 and IQD14 using pairwise transformation analysis.

Supplemental Figure S5. Representative ITC thermogram plots for binding analysis of CML13 to synthetic peptides of IQD14.

Supplemental Figure S6. Overlay protein-protein interaction blots to assess binding of negative control, CML42, to IQD14 *in vitro*.

Supplemental Figure S7. Ca2^+^ titration analysis using a chromophoric chelator and CML13 in the presence or absence of an IQ-domain peptide.

Supplemental Figure S8. Split-luciferase *in planta* protein-interaction assays of the N- and C-lobes of CaM, CML13, CML14 with IQD14.

## List of Supplemental Tables

Supplemental Table S1: Arabidopsis CML protein expression data from public databases.

Supplemental Table S2: Arabidopsis *CML13* and *CML14* transcript expression data from public databases.

Supplemental Table S3: List of oligonucleotide primers used in this study.

Supplemental Table S4: Thermodynamic values from ITC analysis of CaM, CML13, and CML14 binding to synthetic peptides from IQD14 in the presence of ETGA

Supplemental Table S5: Thermodynamic values from ITC analysis of CaM, CML13, and CML14 binding to synthetic peptides from IQD14 in the presence of Ca^2+^

