## Supplementary material for "Arabidopsis calmodulin-like proteins CML13 and CML14 interact with proteins that have IQ domains": Combined Supp Figs and Tables

AtGenExpress eFP: AT1G12310 / CML13

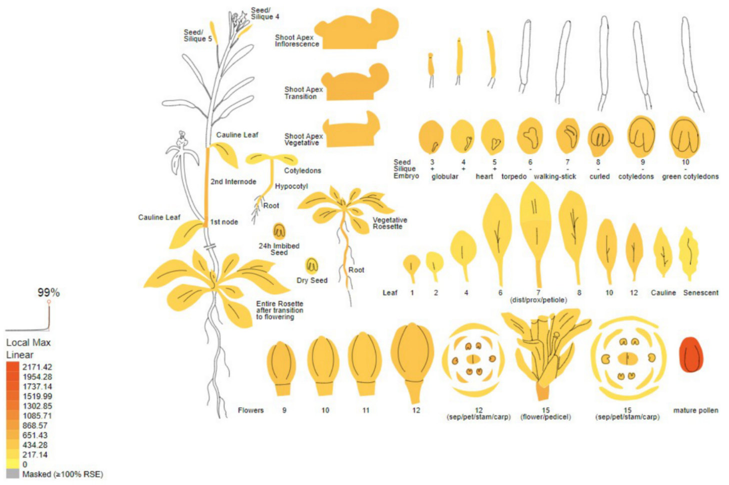


AtGenExpress eFP: AT1G62820 / CML14

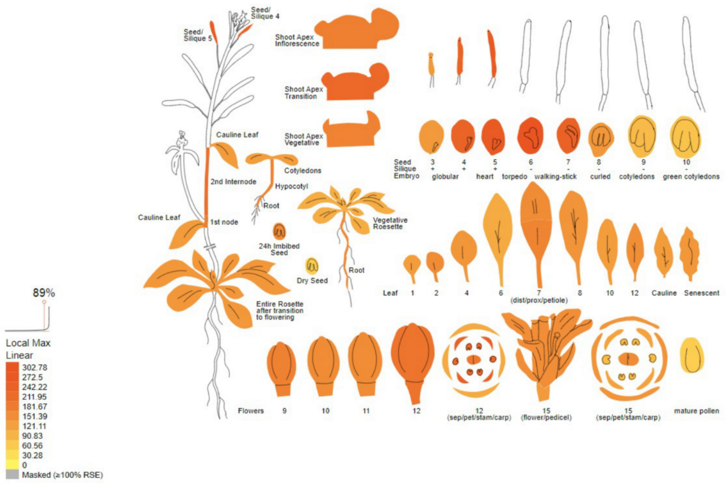


**Supplemental Figure 1.** eFP browser view of *CML13* (upper panel) and *CML14* (lower panel) gene expression during Arabidopsis development. The Arabidopsis eFP Browser is located at bar.utoronto.ca, published in Winter et al. (2007), developed by B. Vinegar, drawn by J. Alls and N. Provart. Data from Gene Expression Map of Arabidopsis Development by Schmid et al. (2005) and the Nambara lab for seed stages. Areas without colouration indicate data was not available.

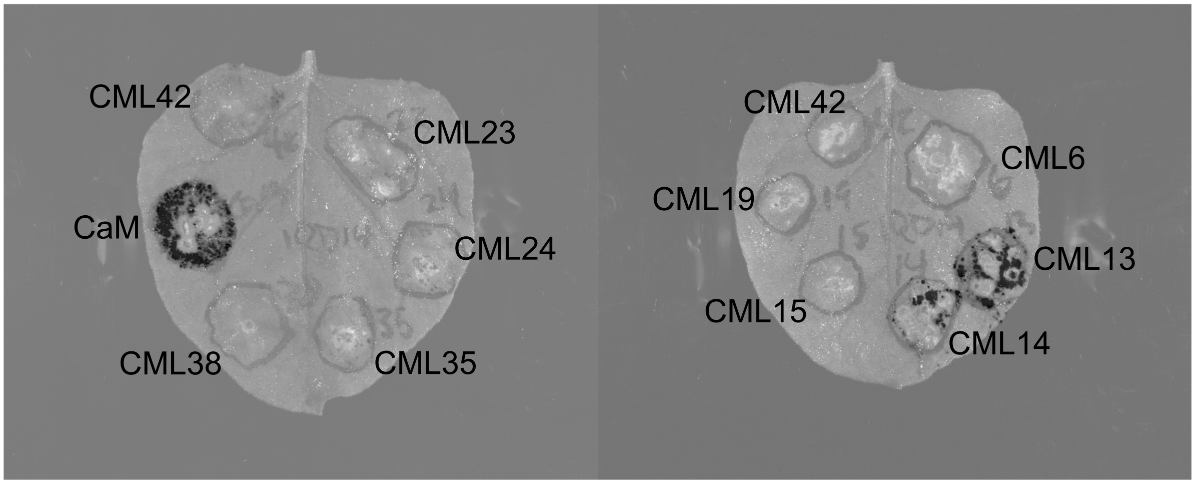


**Supplemental Figure 2**. Whole-leaf images of split-luciferase protein-protein interaction bioluminescent assays using transiently transformed *N. benthamiana*. Representative images are shown for interaction analysis of various CML or CaM baits (as C-Luc fusion proteins) with IQD14 IQ1+2+3 (as N-luc fusion) prey protein. See also Fig.5 legend and Methods for additional details.

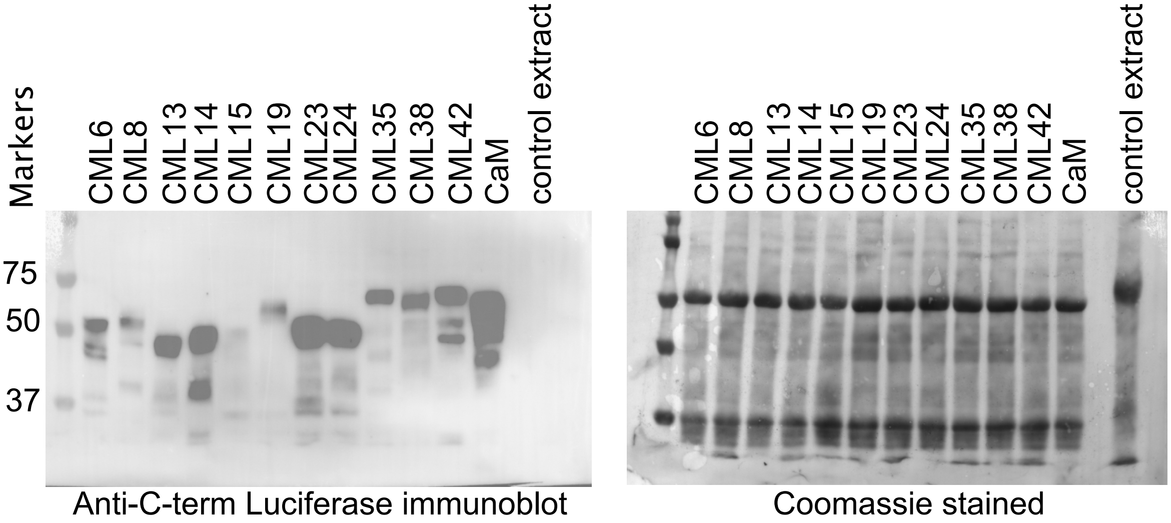


**Supplementary Figure 3**. Immunoblot showing that C-Luc-CML/CaM fusion proteins are expressed in transiently-transformed *N. benthamiana* leaves. Twenty-five µg of crude extracts from transformed leaves (see Methods) were run on SDS-PAGE and subjected to immunoblotting (left panel) with anti-C-terminal luciferase antisera or coomassie stained. Molecular markers are shown on the left (kD).

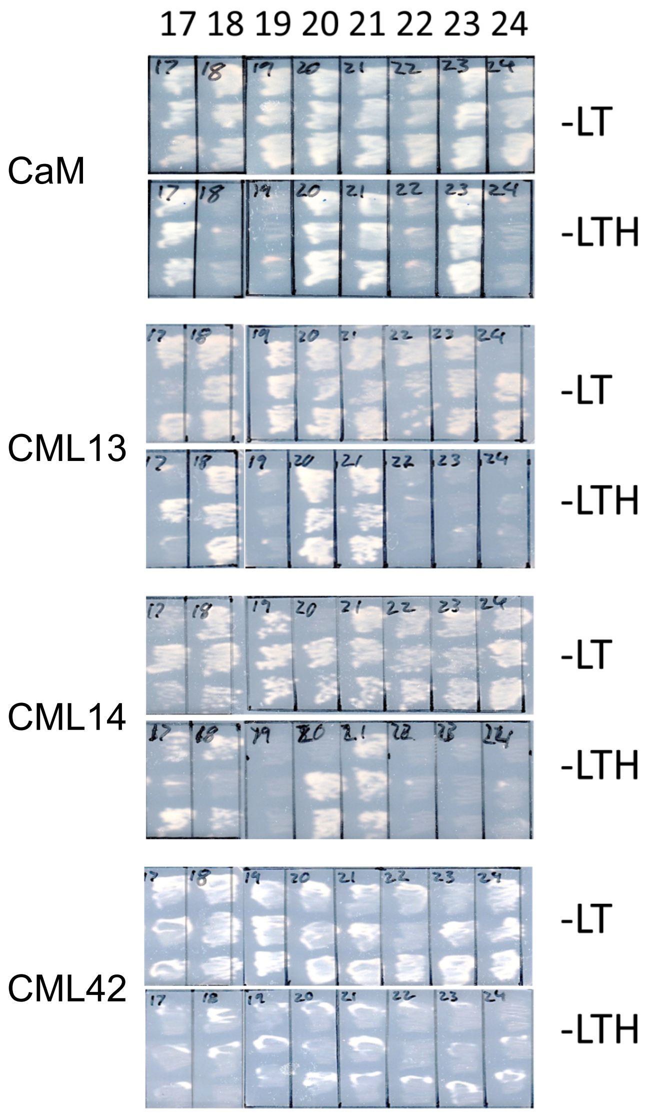


**Supplemental Figure 4:** Delineation of interaction region between CML13/14 and IQ target proteins using yeast two-hybrid pairwise transformation analysis. Bait CMLs or CaM (as indicated) were cotransformed into yeast with respective prey plasmids 17-24 (listed below) and restreaked in triplicate onto -LW (upper panels), -LWH (lower panels) SD media. See Fig. 7 for schematic diagram of IQD14 regions for delineation. Prey plasmids: 17,IQD14-IQ1+2+3; 18, IQD14-IQ2+3; 19, IQD14-IQ3; 20, IQD14-IQ1+2; 21, IQD14-IQ1; 22, IQD14-IQ2; 23, CNGC20 (full-length protein); 24, pGADT7 (empty vector control).

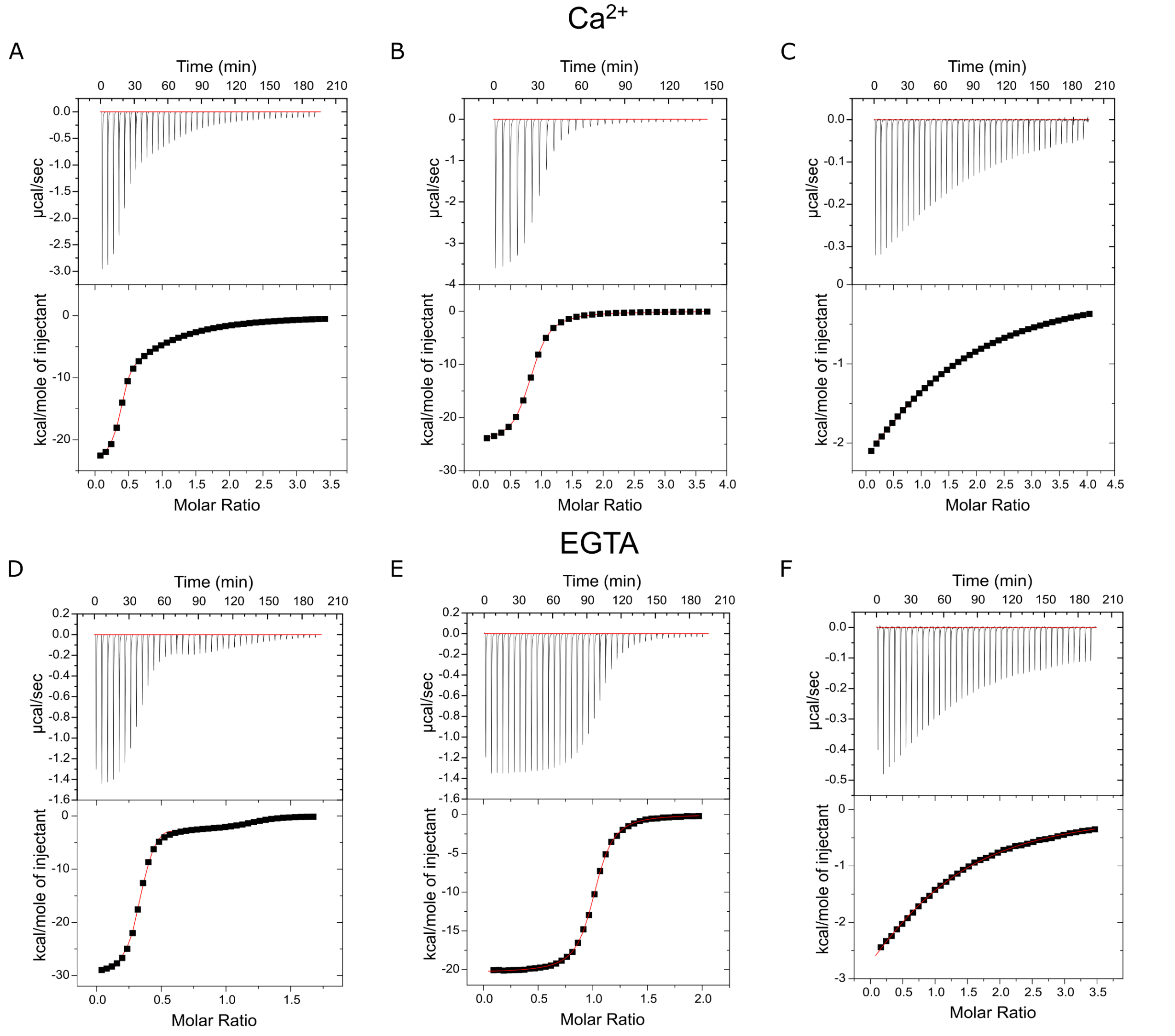


**Supplementary Figure 5**. Representative ITC thermogram plots for binding analysis of CML13 in the presence of 2 mM $CaCl_2$ (A-C) or 2 mM EGTA (D-F) with peptides (A, D) IQD14-IQ1+2, (B,E) IQD14-IQ1, or (C,F) IQD14-IQ2. Titrations were performed as described in the legend for Figure 9. CML13 was used at concentrations from 193-538 µM and IQD14 peptides in the range of 26-35 µM.

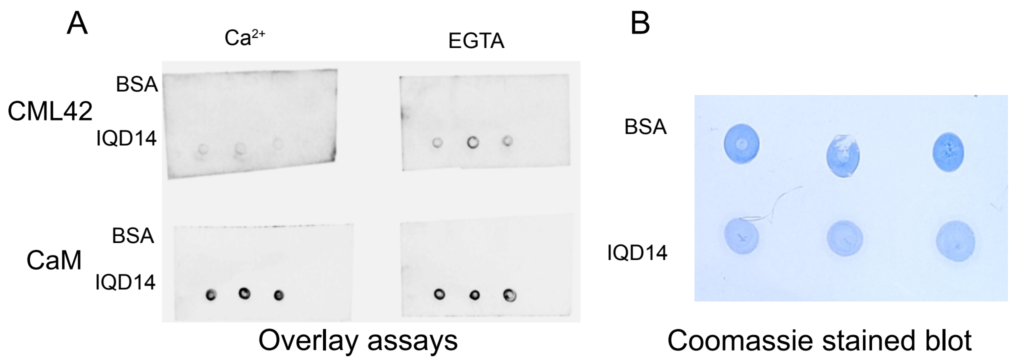


**Supplemental Figure 6.** *In vitro* protein-interaction overlay assays of negative control, CML42, interaction with with IQD14. A) Triplicate samples of pure, recombinant IQD14 (lower spots, 200 ng, corresponding to residues 307-418) or negative control BSA (200 ng, upper spots), were spotted onto nitrocellulose, blocked with 5% casein, and incubated with 200 nM fluorescently-labelled CML42 (upper panels) or positive control, CaM (lower panels), in the presence of 2 mM $CaCl_2$ or 5 mM EGTA as indicated and interaction was detected using a LI-COR infra-red imager. B) A representative Coomassie-stained blot is presented.

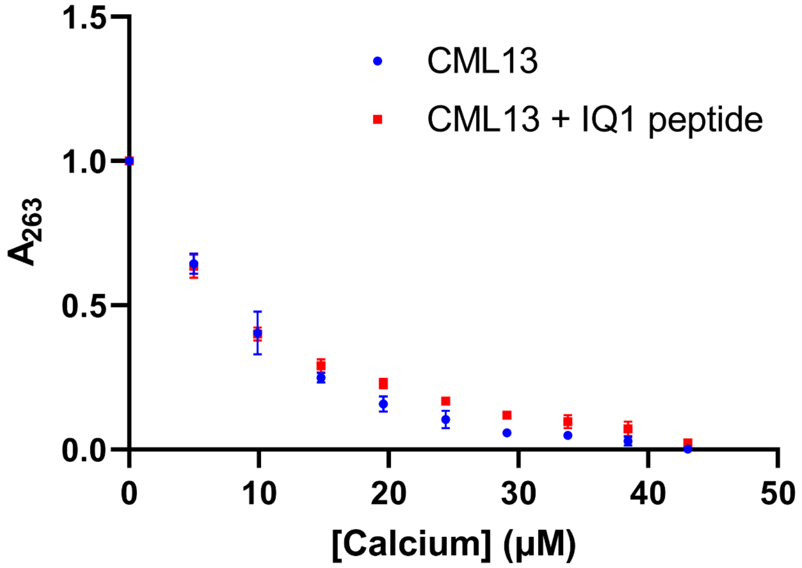


**Supplementary Figure 7**. $Ca^{2+}$ chelator titration assay of CML13. Representative $Ca^{2+}$ titration data for CML13 was obtained by absorption spectroscopy in the presence of the chromophoric chelator 5,5'-$Br_2$-BAPTA at 28 µM. Titration values (means +/- SEM) for CML13 alone (circles) or in the presence of excess IDQ14-IQ1 (IQ1) peptide (squares) circles) are presented based on triplicate assays. The Y-axis was normalized as follows: $Y = (A_{263} - A_{Ca})/(A_0 - A_{Ca})$, where $A_{263}$ is the absorbance at $\lambda$ = 263 nm. Data were analyzed using CaLigator software (Andre et al., 2002).

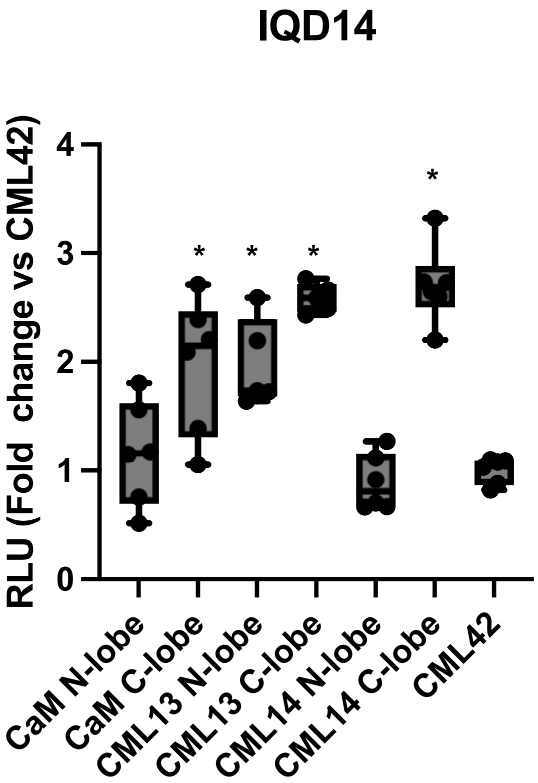


**Supplementary Figure 8**. Interaction of the IQ-domain region of IQD14 with the N- and C-lobes of Arabidopsis CaM, CML13, and CML14. Box plots are presented showing split-luciferase, *in planta* assays to test for binding of IQD14 to respective N- or C-lobes of CML13, CML14, and CaM. *N. benthamiana* leaves were infiltrated with Agrobacterium harbouring respective NLuc-prey and CLuc-bait (CaM, CMLs) vectors and tested for luciferase activity 3 days later, as described in Experimental Procedures. The IQD-region of IQD14 corresponds to residues 306-417. The N-lobes correspond to residues 1-80 for each of CaM, CML13, and CML14. The C-lobes correspond to residues 73-149 for CaM and 72-148 for CML13 and CML14. Boxes contain each data point for 6 technical replicates, means are shown by a horizontal bar, the grey region is the 95% confidence interval, and whiskers extend to maximum and minimum data points. Asterisks indicate a significantly higher signal vs CLuc-CML42 (RLU set to 1) as a negative control bait (pairwise t-test, p-value < 0.05). Data are representative of at least three independent experiments. RLU; relative light units.

**Supplemental Table S1**. Heat map of proteomic data for Arabidopsis CaM and CML expression in various tissues as determined by mass spectrometry analyses and coloured relative to abundance (light green, low; dark green, high) as described in Wang et a., (2021).

| Protein | Calcium sensor | A.thaliana - Leaf, Short da | Leaf, Short | Leaf, SC (Gf | Leaf, SC (Gf | Rosette, En | Rosette, En | Shoot, SC (I | Carpel, SC ( | Cotyledon, | Flower, SC | Flowerbud, | Juvenile lea | Pollen, SC ( | Root, SC (B | Seed, SC (B | Silique, SC | Whole orga | Carpel (Inte | Cotyledon | Flower (Int | Flowerbud | Whole orga | Juvenile lea | Leaf (Integr | Pollen (Inte | Root (Integ | Rosette (Int | Seed (Integ | Shoot (Inte | Silique (Int | Whole orga | Carpel, SC ( | Cotyledon, | Flower, SC | Flowerbud, | Juvenile lea | Leaf, SC (Ba | Pollen, SC ( | Root, SC (B | Seed, SC (B | Shoot, SC ( | Silique, SC | Leaf, SC (Pi | Whole orga | Whole organism, SC (Baerenfaller,science,2008) |
|---|---|---|---|---|---|---|---|---|---|---|---|---|---|---|---|---|---|---|---|---|---|---|---|---|---|---|---|---|---|---|---|---|---|---|---|---|---|---|---|---|---|---|---|---|---|---|
| 3702.AT5G37780.3 | CAM1 | NA | NA | NA | NA | NA | NA | NA | NA | NA | NA | NA | NA | 80.3 | NA | NA | NA | NA | NA | 43.2 | 5.27 | 78.6 | 763 | 145 | 23.2 | 103 | 115 | NA | 49.5 | 33.4 | 134 | 250 | NA | 47.5 | 58 | 94.3 | 174 | 137 | 124 | 218 | 297 | 46.8 | 268 | 69 | 362 | NA |
| 3702.AT2G41110.2 | CAM2 | NA | NA | NA | NA | NA | NA | NA | NA | NA | NA | NA | NA | NA | NA | NA | NA | NA | NA | NA | 0.67 | NA | 1216 | NA | 22 | NA | 8.13 | NA | NA | NA | 9.6 | 391 | NA | NA | 7.39 | NA | NA | NA | NA | 15.4 | NA | NA | 19.2 | 82.2 | 374 | NA |
| 3702.AT3G56800.1 | CAM3 | NA | NA | NA | NA | NA | NA | NA | NA | NA | NA | NA | NA | NA | NA | NA | NA | NA | NA | NA | 0.94 | NA | 1636 | NA | 27.5 | NA | 11.8 | NA | NA | NA | 12.7 | 531 | NA | NA | 10.3 | NA | NA | NA | NA | 22.4 | NA | NA | 25.3 | 103 | 648 | NA |
| 3702.AT1G66410.2 | CAM4 | NA | NA | NA | NA | NA | NA | NA | NA | NA | NA | NA | NA | 93.4 | NA | NA | NA | NA | NA | 49.6 | 6.57 | 93 | 897 | 168 | 25.7 | 121 | 138 | NA | 57 | 40.1 | 159 | 294 | NA | 54.6 | 72.2 | 112 | 202 | 167 | 146 | 261 | 342 | 56.2 | 318 | 74.2 | 434 | NA |
| 3702.AT2G27030.3 | CAM5 | NA | NA | NA | NA | NA | NA | NA | NA | NA | NA | NA | NA | NA | NA | NA | NA | NA | NA | NA | 0.93 | NA | 1620 | NA | 27.2 | NA | 11.6 | NA | NA | NA | 12.5 | 526 | NA | NA | 10.2 | NA | NA | NA | NA | 22.1 | NA | NA | 25 | 102 | 636 | NA |
| 3702.AT3G43810.1 | CAM7 | NA | NA | NA | NA | NA | NA | NA | NA | NA | NA | NA | NA | NA | NA | NA | NA | NA | NA | NA | NA | NA | 1664 | NA | 27.5 | NA | NA | NA | NA | NA | NA | 540 | NA | NA | NA | NA | NA | NA | NA | NA | NA | NA | NA | 103 | 646 | NA |
| 3702.AT3G59450.1 | CML1 | NA | NA | NA | NA | NA | NA | NA | NA | NA | NA | NA | NA | NA | NA | NA | NA | NA | NA | NA | NA | NA | NA | NA | NA | NA | NA | NA | NA | NA | NA | NA | NA | NA | NA | NA | NA | NA | NA | NA | NA | NA | NA | NA | NA | NA |
| 3702.AT4G12860.1 | CML2 | NA | NA | NA | NA | NA | NA | NA | NA | NA | NA | NA | NA | 94.8 | NA | NA | NA | NA | NA | NA | 2.86 | NA | 0.63 | NA | NA | 83.3 | NA | NA | NA | NA | NA | 0.29 | NA | NA | 31.5 | NA | NA | NA | 72.9 | NA | NA | NA | NA | NA | 2.85 | NA |
| 3702.AT3G07490.1 | CML3 | NA | NA | NA | NA | NA | NA | NA | NA | NA | NA | NA | NA | NA | NA | NA | NA | NA | NA | NA | NA | NA | NA | NA | NA | NA | NA | NA | NA | NA | NA | NA | NA | NA | NA | NA | NA | NA | NA | NA | NA | NA | NA | NA | NA | NA |
| 3702.AT3G59440.1 | CML4 | NA | NA | NA | NA | NA | NA | NA | NA | NA | NA | NA | NA | NA | NA | NA | NA | NA | NA | NA | NA | NA | NA | NA | NA | NA | NA | NA | NA | NA | NA | 0.01 | NA | NA | NA | NA | NA | NA | NA | NA | NA | NA | NA | NA | 0.48 | NA |
| 3702.AT2G43290.1 | CML5 | NA | NA | NA | NA | NA | NA | NA | NA | NA | NA | NA | NA | NA | 28.9 | NA | NA | NA | NA | NA | NA | NA | NA | NA | NA | NA | 15.2 | NA | NA | NA | 2.6 | 0.64 | NA | NA | NA | NA | NA | NA | NA | 2.78 | NA | NA | 5.21 | NA | 2.06 | 2.81 |
| 3702.AT4G03290.1 | CML6 | NA | NA | NA | NA | NA | NA | NA | NA | NA | NA | NA | NA | NA | NA | NA | NA | NA | NA | NA | NA | NA | NA | NA | NA | NA | NA | NA | NA | NA | NA | NA | NA | NA | NA | NA | NA | NA | NA | NA | NA | NA | NA | NA | NA | NA |
| 3702.AT1G05990.1 | CML7 | NA | NA | NA | NA | NA | NA | NA | NA | NA | NA | NA | NA | NA | NA | NA | NA | NA | NA | NA | NA | NA | NA | NA | NA | NA | NA | NA | NA | NA | NA | NA | NA | NA | NA | NA | NA | NA | NA | NA | NA | NA | NA | NA | NA | NA |
| 3702.AT4G14640.1 | CML8 | NA | NA | NA | NA | NA | NA | NA | NA | NA | NA | NA | NA | NA | NA | NA | NA | NA | NA | NA | NA | NA | 0.56 | NA | NA | NA | NA | NA | NA | NA | NA | 0.17 | NA | NA | NA | NA | NA | NA | NA | NA | NA | NA | NA | NA | NA | NA |
| 3702.AT3G51920.1 | CML9 | NA | NA | NA | NA | NA | NA | NA | NA | NA | NA | NA | NA | NA | NA | NA | NA | NA | NA | NA | NA | NA | 2.1 | NA | 0.36 | NA | 0.96 | NA | NA | NA | NA | 0.68 | NA | NA | NA | NA | NA | 10.5 | NA | 1.82 | NA | NA | NA | NA | 0.71 | NA |
| 3702.AT2G41090.1 | CML10 | NA | NA | 42.1 | NA | 82.3 | 66.5 | NA | NA | NA | NA | NA | NA | NA | NA | NA | NA | NA | NA | 22.4 | NA | NA | 12.3 | NA | 1.5 | NA | NA | 73 | NA | 18 | NA | 3.89 | NA | 24.6 | NA | NA | NA | 4.96 | NA | NA | NA | 25.2 | NA | NA | 1.88 | NA |
| 3702.AT3G22930.1 | CML11 | NA | NA | NA | NA | NA | NA | NA | NA | NA | NA | NA | NA | NA | NA | NA | NA | NA | NA | NA | NA | NA | NA | NA | NA | NA | 3.27 | NA | NA | NA | NA | 1.28 | NA | NA | NA | NA | NA | NA | NA | 6.22 | NA | NA | NA | NA | 41.2 | NA |
| 3702.AT2G41100.4 | CML12 | 97.6 | 76.5 | 19.6 | 114 | 53.9 | 94.9 | 175 | NA | 144 | 79 | 20.5 | 330 | NA | NA | NA | NA | NA | NA | 61.5 | 73.1 | 20.8 | 226 | 173 | 65.5 | NA | 1.81 | 78 | NA | 154 | NA | 80.9 | NA | 53.2 | 14.3 | 20.9 | 142 | 16.8 | NA | 3.43 | NA | 146 | NA | NA | 96.4 | 35.5 |
| 3702.AT1G12310.1 | CML13 | 37.8 | 31.3 | NA | NA | 195 | 215 | 78.7 | 62.4 | 80.7 | 66.7 | NA | 242 | 450 | 339 | 27.4 | NA | NA | 55.4 | 56.7 | 283 | 95.2 | 262 | 135 | 34.7 | 429 | 448 | 207 | 26.8 | 119 | 990 | 135 | 54 | 54.3 | 2444 | 114 | 114 | 672 | 410 | 547 | 23.9 | 135 | 1979 | NA | 1106 | 93.7 |
| 3702.AT1G62820.1 | CML14 | NA | NA | NA | NA | 90.8 | 45.3 | NA | NA | 80.7 | NA | NA | NA | 58 | 96.8 | NA | NA | NA | NA | 63.8 | 70.6 | 25.4 | 129 | 71.2 | 11 | 47.7 | 120 | 64.1 | NA | 131 | 186 | 67 | NA | 62.1 | 776 | 30.5 | 85.4 | 316 | 38.4 | 140 | NA | 184 | 372 | NA | 602 | 38 |
| 3702.AT1G18530.1 | CML15 | NA | NA | NA | NA | NA | NA | NA | NA | NA | NA | NA | NA | NA | NA | NA | NA | NA | NA | NA | NA | NA | NA | NA | NA | NA | NA | NA | NA | NA | NA | NA | NA | NA | NA | NA | NA | NA | NA | NA | NA | NA | NA | NA | NA | NA |
| 3702.AT3G25600.1 | CML16 | NA | NA | NA | NA | NA | NA | NA | NA | NA | NA | NA | NA | NA | NA | NA | NA | NA | NA | NA | 0.48 | NA | NA | NA | NA | NA | NA | NA | NA | NA | 4.99 | 0.57 | NA | NA | 5.29 | NA | NA | NA | NA | NA | NA | NA | 9.99 | NA | 1.62 | 2.5 |
| 3702.AT1G32250.1 | CML17 | NA | NA | NA | NA | NA | NA | NA | NA | NA | NA | NA | NA | NA | NA | NA | NA | NA | NA | NA | 0.38 | NA | NA | NA | NA | 5.48 | NA | NA | NA | NA | NA | 0.05 | NA | NA | 4.17 | NA | NA | NA | 10.4 | NA | NA | NA | NA | NA | 1.67 | NA |
| 3702.AT3G03000.1 | CML18 | NA | NA | NA | NA | NA | NA | NA | NA | NA | NA | NA | NA | NA | 32.5 | NA | NA | NA | NA | NA | NA | NA | NA | NA | NA | NA | 18 | NA | NA | 8.68 | NA | 0.78 | NA | NA | NA | NA | NA | NA | NA | 5.01 | NA | 12.2 | NA | NA | 3.47 | 3.25 |
| 3702.AT4G37010.2 | CML19 | NA | NA | NA | NA | NA | NA | NA | NA | NA | NA | NA | NA | NA | NA | NA | NA | NA | NA | NA | NA | NA | 2.23 | NA | NA | NA | 7.55 | NA | NA | NA | NA | 0.88 | NA | NA | NA | NA | NA | NA | NA | 14.3 | NA | NA | NA | NA | 5.89 | NA |
| 3702.AT3G50360.1 | CML20 | NA | NA | 70.6 | 37.1 | NA | NA | NA | NA | NA | NA | NA | 75.3 | 21.3 | 201 | NA | NA | NA | NA | 21.8 | 17.8 | 25 | 32.8 | 77.7 | 14.5 | 26.4 | 123 | NA | NA | 44.4 | 9.57 | 20.7 | NA | 24 | 195 | 30 | 78.2 | 17.4 | 30.9 | 53.2 | NA | 62.2 | 19.1 | NA | 93.1 | 36.7 |
| 3702.AT4G26470.1 | CML21 | NA | NA | NA | NA | NA | NA | NA | NA | NA | NA | NA | NA | 37.1 | 36 | NA | NA | NA | NA | NA | NA | NA | NA | NA | NA | 33.7 | 30.1 | NA | NA | NA | NA | 0.83 | NA | NA | NA | NA | NA | NA | 30.7 | 24.8 | NA | NA | NA | NA | 1.42 | 3.8 |
| 3702.AT3G24110.1 | CML22 | NA | NA | NA | 17.8 | NA | NA | NA | NA | NA | NA | NA | NA | NA | NA | NA | NA | NA | NA | NA | NA | NA | NA | NA | 5.61 | NA | 0.55 | NA | NA | NA | NA | 0.01 | NA | NA | NA | NA | NA | NA | NA | 1.04 | NA | NA | NA | NA | 0.48 | NA |
| 3702.AT1G66400.1 | CML23 | NA | NA | NA | NA | NA | NA | NA | NA | NA | NA | NA | NA | NA | NA | NA | NA | NA | NA | NA | 0.79 | NA | NA | NA | NA | NA | NA | NA | NA | NA | NA | 0.7 | NA | NA | 8.68 | NA | NA | NA | NA | NA | NA | NA | NA | NA | 4.87 | 2.66 |
| 3702.AT5G37770.1 | CML24 | NA | NA | NA | NA | NA | NA | NA | NA | NA | NA | NA | NA | NA | 41.4 | NA | NA | NA | NA | NA | 5.16 | 31.8 | 8.48 | 30.1 | 0.16 | NA | 21.7 | NA | NA | NA | NA | 25.1 | NA | NA | 56.8 | 38.2 | 36.2 | 4.56 | NA | 3.97 | NA | NA | NA | NA | 420 | 45.4 |
| 3702.AT1G24620.1 | CML25 | NA | NA | NA | NA | NA | NA | NA | NA | NA | NA | NA | 24.8 | 421 | NA | NA | NA | NA | NA | NA | 6.92 | NA | 1.48 | 12.1 | NA | 410 | 1.17 | NA | NA | NA | NA | 0.87 | NA | NA | 76.1 | NA | 9.5 | NA | 400 | 2.22 | NA | NA | NA | NA | 4.14 | 1.37 |
| 3702.AT1G73630.1 | CML26 | NA | NA | NA | NA | NA | NA | NA | NA | NA | 43.5 | NA | NA | 350 | 31.7 | NA | NA | NA | NA | NA | 51 | 10.4 | 10.9 | NA | 0.29 | 318 | 24.2 | NA | NA | NA | 182 | 6.61 | NA | NA | 126 | 12.5 | NA | 8.4 | 290 | 17.4 | NA | NA | 364 | NA | 32.2 | 10.7 |
| 3702.AT1G18210.1 | CML27 | 115 | 57.7 | 102 | 74.3 | 255 | 170 | NA | 125 | 75.4 | 24.7 | NA | 61.9 | NA | 544 | 97.5 | 191 | NA | 108 | 135 | 185 | 91.2 | 557 | 124 | 65.8 | NA | 585 | 205 | 97.4 | 160 | 2397 | 219 | 105 | 140 | 1787 | 109 | 137 | 436 | NA | 623 | 96.7 | 225 | 4604 | NA | 654 | 118 |
| 3702.AT3G03430.1 | CML28 | NA | NA | NA | NA | NA | NA | NA | NA | NA | NA | NA | NA | NA | NA | NA | NA | NA | NA | NA | 15.4 | 33.1 | 3.67 | NA | NA | 13.1 | NA | NA | NA | NA | NA | 2.03 | NA | NA | 169 | 39.7 | NA | NA | 24.8 | NA | NA | NA | NA | NA | 28.3 | NA |
| 3702.AT5G17480.1 | CML29 | NA | NA | NA | NA | NA | NA | NA | NA | NA | NA | NA | NA | 277 | NA | NA | NA | NA | NA | NA | 23.1 | NA | 4.59 | NA | NA | 265 | NA | NA | NA | NA | NA | 2.38 | NA | NA | 254 | NA | NA | NA | 255 | NA | NA | NA | NA | NA | 30.3 | NA |
| 3702.AT2G15680.1 | CML30 | NA | NA | NA | NA | NA | NA | NA | NA | NA | NA | NA | NA | NA | NA | NA | NA | NA | NA | NA | NA | NA | NA | NA | NA | NA | NA | NA | NA | NA | NA | 0.01 | NA | NA | NA | NA | NA | NA | NA | NA | NA | NA | NA | NA | 0.46 | NA |
| 3702.AT2G36180.1 | CML31 | NA | NA | 28.5 | NA | NA | NA | NA | NA | NA | NA | NA | NA | NA | NA | NA | NA | NA | NA | NA | NA | NA | NA | NA | 0.9 | NA | NA | NA | NA | NA | NA | NA | NA | NA | NA | NA | NA | NA | NA | NA | NA | NA | NA | NA | NA | NA |
| 3702.AT5G17470.1 | CML32 | NA | NA | NA | NA | NA | NA | NA | NA | NA | NA | NA | NA | NA | NA | NA | NA | NA | NA | NA | NA | NA | NA | NA | NA | NA | NA | NA | NA | NA | NA | NA | NA | NA | NA | NA | NA | NA | NA | NA | NA | NA | NA | NA | NA | NA |
| 3702.AT3G03400.1 | CML33 | NA | NA | NA | NA | NA | NA | NA | NA | NA | NA | NA | NA | NA | NA | NA | NA | NA | NA | NA | NA | NA | NA | NA | NA | NA | NA | NA | NA | NA | NA | NA | NA | NA | NA | NA | NA | NA | NA | NA | NA | NA | NA | NA | NA | NA |
| 3702.AT3G03410.1 | CML34 | NA | NA | NA | NA | NA | NA | NA | NA | NA | NA | NA | NA | NA | NA | NA | NA | NA | NA | NA | NA | NA | NA | NA | NA | NA | NA | NA | NA | NA | NA | NA | NA | NA | NA | NA | NA | NA | NA | NA | NA | NA | NA | NA | NA | NA |
| 3702.AT2G41410.1 | CML35 | NA | NA | NA | NA | NA | NA | NA | NA | 35.2 | NA | NA | NA | NA | 73.4 | NA | NA | NA | NA | 16.9 | 2.43 | NA | 15.1 | NA | 0.37 | NA | 41.9 | NA | NA | NA | 4.16 | 8.09 | NA | 15.1 | 26.7 | NA | NA | 10.5 | NA | 13.5 | NA | NA | 8.33 | NA | 35.5 | 10.9 |
| 3702.AT3G10190.1 | CML36 | NA | NA | NA | NA | NA | NA | NA | 104 | NA | NA | NA | NA | NA | NA | NA | NA | NA | 93 | NA | NA | 15.2 | NA | NA | 0.13 | NA | NA | NA | NA | NA | NA | 1.21 | 90.7 | NA | NA | 18.2 | NA | 3.75 | NA | NA | NA | NA | NA | NA | 4.9 | 5.14 |
| 3702.AT5G42380.1 | CML37 | NA | NA | NA | 19.90 | NA | NA | NA | NA | NA | NA | NA | NA | NA | NA | NA | NA | NA | NA | NA | NA | NA | 22.5 | NA | 6.29 | NA | NA | NA | NA | NA | NA | 7.43 | NA | NA | NA | NA | NA | NA | NA | NA | NA | NA | NA | NA | 0.24 | 1.91 |
| 3702.AT1G76650.1 | CML38 | NA | NA | NA | NA | NA | NA | NA | NA | NA | NA | NA | NA | NA | NA | NA | NA | NA | NA | NA | NA | NA | 5.73 | NA | NA | NA | NA | NA | NA | NA | NA | 2.36 | NA | NA | NA | NA | NA | NA | NA | NA | NA | NA | NA | NA | 5.35 | 1.94 |
| 3702.AT1G76640.1 | CML39 | NA | NA | NA | NA | NA | NA | NA | NA | NA | NA | NA | NA | NA | NA | NA | NA | NA | NA | NA | NA | NA | NA | NA | NA | NA | NA | NA | NA | NA | NA | 0.01 | NA | NA | NA | NA | NA | NA | NA | NA | NA | NA | NA | NA | 0.33 | NA |
| 3702.AT3G01830.1 | CML40 | NA | NA | NA | NA | NA | NA | NA | NA | NA | NA | NA | NA | NA | NA | NA | NA | NA | NA | NA | 3.43 | NA | 11.8 | NA | NA | NA | NA | NA | NA | NA | NA | 4.02 | NA | NA | 37.7 | NA | NA | NA | NA | NA | NA | NA | NA | NA | 10.6 | NA |
| 3702.AT3G50770.1 | CML41 | NA | NA | NA | NA | NA | NA | NA | NA | NA | NA | NA | NA | NA | NA | NA | NA | NA | NA | NA | NA | NA | 1.57 | NA | NA | NA | NA | NA | NA | NA | NA | 0.49 | NA | NA | NA | NA | NA | NA | NA | NA | NA | NA | NA | NA | NA | NA |
| 3702.AT4G20780.1 | CML42 | NA | NA | NA | NA | NA | NA | NA | NA | NA | NA | NA | NA | 627 | 430 | NA | NA | NA | NA | NA | 16 | 110 | 14.7 | NA | 0.38 | 564 | 222 | NA | NA | 12.9 | NA | 13.5 | NA | NA | 176 | 132 | NA | 10.9 | 507 | 35 | NA | 18.1 | NA | NA | 19.7 | 40.5 |
| 3702.AT5G44460.1 | CML43 | NA | NA | NA | NA | NA | NA | NA | NA | NA | NA | NA | NA | NA | 51.4 | NA | NA | NA | NA | NA | NA | NA | 2.9 | NA | NA | NA | 26.9 | NA | NA | NA | NA | 2.06 | NA | NA | NA | NA | NA | NA | NA | 4.91 | NA | NA | NA | NA | 4.01 | 4.97 |
| 3702.AT1G21550.1 | CML44 | NA | NA | NA | NA | NA | NA | NA | NA | NA | NA | NA | NA | NA | NA | NA | NA | NA | NA | NA | NA | NA | NA | NA | NA | NA | NA | NA | NA | NA | NA | NA | NA | NA | NA | NA | NA | NA | NA | NA | NA | NA | NA | NA | NA | NA |
| 3702.AT3G29000.1 | CML45 | NA | NA | NA | NA | NA | NA | NA | NA | NA | NA | NA | NA | NA | NA | NA | NA | NA | NA | NA | NA | NA | 0.86 | NA | NA | NA | NA | NA | NA | NA | NA | 0.27 | NA | NA | NA | NA | NA | NA | NA | NA | NA | NA | NA | NA | NA | NA |
| 3702.AT5G39670.1 | CML46 | NA | NA | NA | NA | NA | NA | NA | NA | NA | NA | NA | NA | NA | NA | NA | NA | NA | NA | NA | NA | NA | NA | NA | NA | NA | NA | NA | NA | NA | NA | 0.02 | NA | NA | NA | NA | NA | NA | NA | NA | NA | NA | NA | NA | 0.59 | NA |
| 3702.AT3G47480.1 | CML47 | NA | NA | NA | NA | NA | NA | NA | NA | NA | NA | NA | NA | NA | NA | NA | NA | NA | NA | NA | NA | NA | NA | NA | NA | NA | NA | NA | NA | NA | NA | NA | NA | NA | NA | NA | NA | NA | NA | NA | NA | NA | NA | NA | NA | NA |
| 3702.AT2G27480.1 | CML48 | NA | NA | NA | NA | NA | NA | NA | NA | NA | NA | NA | NA | NA | NA | NA | NA | NA | NA | NA | NA | NA | NA | NA | NA | NA | NA | NA | NA | NA | NA | NA | NA | NA | NA | NA | NA | NA | NA | NA | NA | NA | NA | NA | NA | NA |
| 3702.AT3G10300.3 | CML49 | NA | NA | NA | NA | NA | NA | NA | 75.9 | NA | 104 | 45.3 | NA | NA | 44.3 | 99.9 | 121 | NA | 70.1 | NA | 98.5 | 95.9 | 33.8 | NA | 0.65 | 6.7 | 25.3 | NA | 97.9 | NA | 81.5 | 28.7 | 69 | NA | 47.4 | 106 | NA | 18.8 | 12.7 | 8.13 | 88.1 | NA | 42.3 | NA | 50.4 | 80.4 |
| 3702.AT5G04170.1 | CML50 | NA | NA | NA | NA | NA | 26.8 | NA | NA | NA | NA | 40.1 | NA | NA | 111 | 38.6 | 10.2 | NA | NA | NA | NA | 24.3 | 11.6 | NA | 3.49 | NA | 119 | 15.8 | 38 | NA | 7.34 | 8.76 | NA | NA | NA | 21.2 | NA | 101 | NA | 127 | 34.8 | NA | 4.53 | NA | 23 | 21.5 |

**Supplementary Table 2**: Absolute developmental transcript expression data for *CML13* and *CML14* sourced from the Arabidopsis eFP expression browser (Winter et al., 2007).

| Locus | CML | | Dry seed | Imbibed seed, 24 h | 1st Node | Flower Stage 12, Stamens | Cauline Leaf | Cotyledon | Root | Entire Rosette After Transition to Flowering | Flower Stage 9 | Flower Stage 10/11 |
|---|---|---|---|---|---|---|---|---|---|---|---|---|
| At1g12310 | 13 | Expression Level | 226.94 | 488.86 | 703.01 | 730.9 | 245.01 | 182.73 | 641.81 | 326.4 | 486.6 | 410.04 |
| | | Standard Deviation | 16.94 | 7.11 | 21.59 | 49.9 | 7.65 | 12.54 | 12.89 | 10.06 | 35.28 | 25.44 |
| At1g62820 | 14 | Expression Level | 52.67 | 154.72 | 176.31 | 219.13 | 127.48 | 124.71 | 156.25 | 133.96 | 171.73 | 154.8 |
| | | Standard Deviation | 6.19 | 38.4 | 21.78 | 9.72 | 24.99 | 6.74 | 15.43 | 12.7 | 10.43 | 16.71 |

| Locus | CML | | Flower Stage 12 | Flower Stage 15 | Flower Stage 12, Carpels | Flower Stage 12, Petals | Flower Stage 12, Sepals | Flower Stage 15, Carpels | Flower Stage 15, Petals | Flower Stage 15, Sepals | Flower Stage 15, Stamen | Flowers Stage 15, Pedicels | Leaf 1 + 2 | Leaf 7, Petiole |
|---|---|---|---|---|---|---|---|---|---|---|---|---|---|---|
| At1g12310 | 13 | Expression Level | 469.55 | 320.65 | 468.83 | 502.88 | 290.88 | 452.25 | 276.58 | 195.06 | 432.18 | 724.96 | 338.4 | 442.73 |
| | | Standard Deviation | 33.77 | 26.24 | 63.53 | 19.48 | 14.79 | 8.3 | 2.02 | 5.58 | 5.57 | 51.03 | 22.02 | 14.99 |
| At1g62820 | 14 | Expression Level | 196 | 148.45 | 188.35 | 243.03 | 104.8 | 154.65 | 116.9 | 138.21 | 78.11 | 129.79 | 125.01 | 160 |
| | | Standard Deviation | 18.7 | 14.89 | 13.78 | 27.88 | 5.48 | 18.08 | 17.5 | 17.74 | 13.47 | 12.22 | 9.99 | 7.46 |

| Locus | CML | | Leaf 7, Distal Half | Leaf 7, Proximal Half | Hypocotyl | Root | Rosette Leaf 2 | Rosette Leaf 4 | Rosette Leaf 6 | Rosette Leaf 8 | Rosette Leaf 10 | Rosette Leaf 12 | Senescing Leaf | Shoot Apex, Inflorescence | Shoot Apex, Transition |
|---|---|---|---|---|---|---|---|---|---|---|---|---|---|---|---|
| At1g12310 | 13 | Expression Level | 229.88 | 300.14 | 367.26 | 578.61 | 174.63 | 234 | 267.78 | 308.2 | 378.25 | 431 | 149.18 | 539.73 | 604.31 |
| | | Standard Deviation | 16.15 | 2.78 | 11.77 | 31.97 | 12.74 | 4.85 | 11.26 | 10.13 | 27.53 | 23.87 | 3.39 | 3.38 | 19.21 |
| At1g62820 | 14 | Expression Level | 154.06 | 163.4 | 168.58 | 206.46 | 152.2 | 136.1 | 105.31 | 137.5 | 125.25 | 147.48 | 132.71 | 175.71 | 221.63 |
| | | Standard Deviation | 23.06 | 19.04 | 6.12 | 12.3 | 9.23 | 12.63 | 10.23 | 4.49 | 13.49 | 13.66 | 10.99 | 31.3 | 5.45 |

| Locus | CML | | Shoot Apex, Vegetative | Stem, 2nd Internode | Mature Pollen | Seeds Stage 3 w/ Siliques | Seeds Stage 4 w/ Siliques | Seeds Stage 5 w/ Siliques | Seeds Stage 6 w/o Siliques | Seeds Stage 7 w/o Siliques | Seeds Stage 8 w/o Siliques | Seeds Stage 9 w/o Siliques | Seeds Stage 10 w/o Siliques | Vegetative Rosette |
|---|---|---|---|---|---|---|---|---|---|---|---|---|---|---|
| At1g12310 | 13 | Expression Level | 505.53 | 609.31 | 2171.41 | 522.26 | 341.71 | 355.25 | 433.58 | 472.03 | 518.43 | 455.15 | 459.73 | 386.8 |
| | | Standard Deviation | 22.06 | 23.9 | 138.67 | 6.68 | 5.85 | 10.92 | 10.98 | 20.09 | 18.59 | 19.65 | 27.05 | 15.37 |
| At1g62820 | 14 | Expression Level | 167.8 | 205.68 | 55.01 | 130.81 | 233.26 | 263.51 | 302.78 | 237.73 | 132.6 | 80.14 | 68.64 | 101.48 |
| | | Standard Deviation | 8.57 | 22.19 | 23.83 | 23.65 | 13.51 | 9.41 | 12.76 | 28.18 | 8.77 | 8.07 | 9.41 | 11.09 |

**Table S2**: List of oligonucleotide primer sequences.

| Primer name | Primer sequence (5'->3') | Notes |
|---|---|---|
| Nluc CNGC20 F | GGCCGGATCCATGCTGAGTGTAGCCGATCTGG | for split luciferase cloning CNGC20 |
| Nluc CNGC20 R | GGCCGGTCGACAAGGCTATAACTAGACTGAGG | for split luciferase cloning CNGC20 |
| Nluc IQM1 F | GGCCGGATCCATGGTCACCGAGCTTGATGCAGC | for split luciferase cloning IQM1 |
| Nluc IQM1 R | GGCCGGTCGACCTCAACAACGACCGCACAATCTG | for split luciferase cloning IQM1 |
| Nluc GAD1 CaMBD F | GGCCGGATCCATGAACAGCGATAACTTGATGG | for split luciferase cloning GAD1 |
| Nluc GAD1 CaMBD R | GGCCGGTCGACGCAGATACCACTCGTCTTCTTCC | for split luciferase cloning GAD1 |
| Cluc CML24 F | GGCCGGTACCATGTCATCGAAGAACGGAG | for split luciferase cloning CML24 |
| Cluc CML24 R | GGCCGGATCCTCAAGCACCACCACCATTACTC | for split luciferase cloning CML24 |
| Cluc CML30 F | GGCCGGTACCATGTCAAACGTGAGTTTTCTTG | for split luciferase cloning CML30 |
| Cluc CML30 R | GGCCGGATCCTTAGACATTGTTGGAAGACATC | for split luciferase cloning CML30 |
| Cluc CML35 F | GGCCGGTACCATGAAGCTCGCCGCTAGCC | for split luciferase cloning CML35 |
| Cluc CML35 R | GGCCGGATCCCTAATGATGATGATCATTCATCGC | for split luciferase cloning CML35 |
| Cluc CML38 F | GGCCGGTACCATGAAGAATAATACTCAACCTC | for split luciferase cloning CML38 |
| Cluc CML38 R | GGCCGGATCCTTAGCGCATCATAAGAGCAAAC | for split luciferase cloning CML38 |
| Cluc CML6 F | GGCCGGTACCATGGATTCCACGGAGCTGAAC | for split luciferase cloning CML6 |
| Cluc CML6 R | GGCCGGATCCTCAGCTCAAAGAGCTAAAAAAGC | for split luciferase cloning CML6 |
| Cluc CML8 F | GGCCGGTACCATGGAAGAAACAGCACTGAC | for split luciferase cloning CML8 |
| Cluc CML8 R | GGCCGGATCCTCAGTCAATGTTGATCATCATC | for split luciferase cloning CML8 |
| Cluc CML15 F | GGCCGGTACCATGGAGGATCAGATAAGACAAC | for split luciferase cloning CML15 |
| Cluc CML15 R | GGCCGGATCCTCACGAATTAATTTTTAATCCAAAG | for split luciferase cloning CML15 |
| Cluc CML19 F | GGCCGGTACCATGGCGAATTACATGTCGG | for split luciferase cloning CML19 |
| Cluc CML19 R | GGCCGGATCCTTAGCCGTAAGAGGTTCTCTTC | for split luciferase cloning CML19 |
| Cluc CML23 F | GGCCGGTACCATGTCGAAGAACGTTTCGAG | for split luciferase cloning CML23 |
| Cluc CML23 R | GGCCGGATCCTCAAGCACTACCATTAATCATC | for split luciferase cloning CML23 |
| HT69 LP | GCAAACCCACTCAAATTATCC | CML13 t-DNA genotyping |
| HT70 RP | CATTTCAAGATCGAAAGAGCG | CML13 t-DNA genotyping |
| HT71 LP | TGTTTCCGTGTAACTTCTGGG | CML14 t-DNA genotyping |
| HT72 RP | TTGTTCTTGACGACGACTGTG | CML14 t-DNA genotyping |
| HT63 Fp | CCGGCCAAGCTTGTTTGGGTTTTAGCTTAAAGTTTGC | CML13 promotor cloning for GUS analysis |
| HT64 Rp | GGCCGGATTCTTTTCGATTCTTCCGGGT | CML13 promotor cloning for GUS analysis |
| HT67 Fp | | CML14 promotor cloning for GUS analysis |
| HT101 Rp | GGCCGGCCATGGCTTCGATTCGGTTCTTACGG | CML14 promotor cloning for GUS analysis |
| HT205 Fp | GGCCGGGGTACCATGGTGAAGAAAGGAAGTTGG | for IQD14 cloning into pRTL2\|mcherry |
| HT206 Rp | GGCCGGTCTAGATCACACAAATCTGTTAAATTCC | for IQD14 cloning into pRTL2\|mcherry |
| HT207 Fp | GGCCGGTCGACATGGCGGATCAGCTTACAG | for cloning CaM into pRTL2\|2xGFP |
| HT208 Rp | GGCCGGAATTCTCACTTGGCCATCATGACC | for cloning CaM into pRTL2\|2xGFP |

| | | |
|---|---|---|
| HT119 Fp | GGCCGGAAGCTTATGGGGAAAGATGGTC | for cloning CML13 into pRTL2\|2xGFP |
| HT120 Rp | GGCCGGGAATTCTCACTTAGCAACCATCC | for cloning CML13 into pRTL2\|2xGFP |
| HT121 Fp | GGCCGGAAGCTTATGAGCAAGGATGGTTTGAG | for cloning CML14 into pRTL2\|2xGFP |
| HT122 Rp | GGCCGGGAATTCTTACTTAGCAACCATTCTAGC | for cloning CML14 into pRTL2\|2xGFP |
| HT81 Fp | GGCCGGTACCATGGAGTACCCATACGACGTACC | HA tag specific, for cloning IQD14, Myosin VIII-B and CAMTA4 into pCAMBIA1300\|NLuc and CML13, CML14, and CaM into pCAMBIA1300\|CLuc |
| HT123 Rp | GGCCGGGTCGACCACAAATCTGTTAAATTCCTTTCTC | for cloning IQD14 into pCAMBIA1300\|NLuc |
| HT124 Rp | GGCCGGGTCGACATAACTTTTCTTGCACCACCAATTCC | for cloning Myosin VIII-B into pCAMBIA1300\|NLuc |
| HT125 Rp | GGCCGGGTCGACAGGCAATGAAAACAAAGTGTCATATTCC | for cloning CAMTA4 into pCAMBIA1300\|NLuc |
| HT162 Fp | GGCCGGAGATCTGATGGGGAAAGATGGTCTGAGC | for cloning CML13 into pCambia1300\|CLuc |
| HT80 Rp | GGCCGTCGACTCACTTAGCAACCATCCTTGC | for cloning CML13 into pCambia1300\|CLuc |
| HT82 Rp | GGCCGTCGACTCAAGAAGAAGGGATGACAACAG | for cloning CaM into pCambia1300\|CLuc |
| HT83 Rp | GGCCGTCGACTCACTTGGCCATCATGACC | for cloning CML42 into pCambia1300\|CLuc |
| HT6 Rp | GGCCGGGTCGACTTACTTAGCAACCATTCTAGCAATAAAATC | for cloning CML14 into pCambia1300\|CLuc |
| HT3 Fp | GGCCGGCATATGGGGAAAGATGGTC | for cloning CML13 into pGBKT7 |
| HT4 Rp | GGCCGGGTCGACTCACTTAGCAACCATCC | for cloning CML13 into pGBKT7 |
| HT5 Fp | GGCCGGCATATGAGCAAGGATGGTTTGAGC | for cloning CML14 into pGBKT7 |
| HT6 Rp | GGCCGGGTCGACTTACTTAGCAACCATTCTAGCAATAAAATC | for cloning CML14 into pGBKT7 and pET30a |
| HT173 Fp | GGCCGGAATTCATGGCGGATCAGCTTACAG | for cloning CaM into pGBKT7 |
| HT174 Rp | GGCCGGTCGACTCACTTGGCCATCATGACCTTG | for cloning CaM into pGBKT7 |
| HT110 Fp | GGCCGGGAATTCCAGCGGCGGGAGA | for cloning Y2H delineation constructs IQD14 IQ67, IQ1+2, and IQ1 |
| HT111 Rp | GGCCGGGTCGACTCATGTCAGCACACTGTCATCC | for cloning Y2H delineation constructs IQD14 IQ67, IQ2+3, and IQ3 |
| HT112 Fp | GGCCGGGAATTCTACATGGCAAGGAAGAGTTTC | for cloning Y2H delineation constructs IQD14 IQ2+3, and IQ2 |
| HT113 Fp | GGCCGGGAATTCAGCGTGAAGCGGCAG | for cloning Y2H delineation construct IQD14 IQ3 |
| HT117 Rp | GGCCGGGTCGACTCAAACACGGACCACTTGCTG | for cloning Y2H delineation constructs IQD14 IQ1+2, and IQ2 |
| HT154 Rp | GGCCGGGTCGACTCACACCACACCTTGAAGTC | for cloning Y2H delineation construct IQD14 IQ1 |

| | | |
|---|---|---|
| HT146 Fp | GGCCGGCATATGATGGCTTCCCACAAC | for cloning CNGC20 into pGADT7 |
| HT147 Rp | GGCCGGGAATTCCTAAAGGCTATAACTAGACTG | for cloning CNGC20 into pGADT7 |
| HT140 Fp | GGCCGGCATATGGATGGGGAAAGATGGTC | for cloning CML13 into pET16b |
| HT141 Rp | GGCCGGCTCGAGTCACTTAGCAACCATCC | for cloning CML13 into pET16b |
| HT59 Fp | GGCCGGAATTCATGAGCAAGGATGGTTTGAGC | for cloning CML14 into pET30a |

**Table S4:** Summary of Raw ITC data obtained for CaM, CML13, and CML14 binding to IQD14 peptides in 10 mM Tris-Cl pH 7.5 and 2 mM EGTA. N, Stoichiometry; $K_a$, Association Constant ($M^{-1}$); ΔH, Enthalpy (cal $mol^{-1}$); ΔS, Entropy (cal $mol^{-1}$ $K^{-1}$).

| EGTA | | | |
|---|---|---|---|
| | IQD14-IQ1+2 | IQD14-IQ1 | IQD14-IQ2 |
| **CaM** | | | |
| $N_1$ | 0.438 | 0.985 | 0.240 |
| $K_{a1}$ | 1.00E9 ± 1.6 E8 | 1.43E8 ± 1.43E7 | 4.79 E6 ± 4.45 E5 |
| $\Delta H_1$ | - 3.29 E4 ± 108 | -3.288 E4 ± 78.28 | -2382 ± 441 |
| $\Delta S_1$ | -67.6 | -71.1 | 22.7 |
| $N_2$ | 0.415 | | 0.377 |
| $K_{a2}$ | 2.64 E6 ± 3.98 E5 | | 7.55 E5 ± 1.61 E4 |
| $\Delta H_2$ | -9744 ± 230 | | -1.511 E4 ± 288 |
| $\Delta S_2$ | -2.76 | | -23.0 |
| **CML13** | | | |
| $N_1$ | 0.316 | 0.999 | 1.02 |
| $K_{a1}$ | 2.87E8 ± 1.62 E8 | 4.5E6 ± 1.86 E5 | 1.81 E4 ± 851 |
| $\Delta H_1$ | - 2.981 E4 ± 269 | -2.036 E4 ± 62.31 | -7855 ± 562.1 |
| $\Delta S_1$ | -59.6 | -36.7 | -6.43 |
| $N_2$ | 0.881 | | |
| $K_{a2}$ | 2.91 E6 ± 1.69 E6 | | |
| $\Delta H_2$ | -2299 ± 187 | | |
| $\Delta S_2$ | 22 | | |
| **CML14** | | | |
| $N_1$ | 0.323 | 0.810 | 0.996 |
| $K_{a1}$ | 8.67 E7 ± 2.07 E7 | 8.65 E6 ± 2.07 E5 | 3.16 E5 ± 4.69 E3 |
| $\Delta H_1$ | - 2.333 E4 ± 341 | -2.385 E4 ± 36.88 | -6398 ± 19.18 |
| $\Delta S_1$ | -40.6 | -46.9 | -4.06 |
| $N_2$ | 0.657 | | |
| $K_{a2}$ | 1.12 E6 ± 2.2 E5 | | |
| $\Delta H_2$ | -9382 ± 388 | | |
| $\Delta S_2$ | -3.27 | | |

**Table S5 :** Summary of Raw ITC data obtained for CaM, CML13, and CML14 binding to IQD14 peptides in 10 mM Tris-Cl pH 7.5 and 2 mM $CaCl_2$. N, Stoichiometry; $K_a$, Association Constant ($M^{-1}$); ΔH, Enthalpy (cal mol−1); ΔS, Entropy (cal mol−1 K−1).

| Calcium | | | |
|---|---|---|---|
| | IQD14-IQ1+2 | IQD14-IQ1 | IQD14-IQ2 |
| **CaM** | | | |
| $N_1$ | 0.695 | 0.329 | 0.479 |
| $K_{a1}$ | 8.44E8 ± 0.00267 | 1.09 E9 ± 7.48 E8 | 3.12 E8 ± 1.23 E8 |
| $\Delta H_1$ | - 1.305 E4 ± 16 | -3.559 E4 ± 730 | -15540 ± 378 |
| $\Delta S_1$ | -2.22 | -76.0 | -12.4 |
| $N_2$ | 0.260 | 0.378 | 0.513 |
| $K_{a2}$ | 2.75 E6 ± 1.07 E5 | 1.16 E8 ± 8.70 E7 | 4.56 E7 ± 2.06 E7 |
| $\Delta H_2$ | -10370 ± 97.2 | -1556 ± 713 | -707.5 ± 416 |
| $\Delta S_2$ | -4.73 | 31.8 | 32.7 |
| **CML13** | | | |
| $N_1$ | 0.348 | 0.801 | 1.16 |
| $K_{a1}$ | 4.6 E6 ± 0.00462 | 1.07 E6 ± 1.92 E4 | 1.74 E4 ± 489 |
| $\Delta H_1$ | - 2.343 E4 ± 188 | -2.496 E4 ± 59.84 | -5683 ± 216 |
| $\Delta S_1$ | -46.8 | -54.7 | 0.656 |
| $N_2$ | 0.725 | | |
| $K_{a2}$ | 6.19 E4 ± 6.93 E3 | | |
| $\Delta H_2$ | -1.339 E4 ± 1770 | | |
| $\Delta S_2$ | -22.3 | | |
| **CML14** | | | |
| $N_1$ | 0.328 | 0.825 | 0.882 |
| $K_{a1}$ | 3.64 E7 ± 3.26E6 | 4.13 E6 ± 5.40 E4 | 1.91 E5 ± 2.5 E3 |
| $\Delta H_1$ | - 2.677 E4 ± 189 | -2.178 E4 ± 23.57 | -4550 ± 17.31 |
| $\Delta S_1$ | -53.7 | -41.6 | 9.15 |
| $N_2$ | 0.668 | | |
| $K_{a2}$ | 8.99 E5 ± 8.05 E4 | | |
| $\Delta H_2$ | -8356 ± 216 | | |
| $\Delta S_2$ | -0.323 | | |
